# MGMM: An R Package for fitting Gaussian Mixture Models on Incomplete Data

**DOI:** 10.1101/2019.12.20.884551

**Authors:** Zachary R. McCaw, Hanna Julienne, Hugues Aschard

**Author notes:** These authors contributed equally to this work (Zachary R. McCaw, Hanna Julienne). **Contact:** Zachary R. McCaw, Hanna Julienne.

## Abstract

Although missing data are prevalent in applications, existing implementations of Gaussian mixture models (GMMs) require complete data. Standard practice is to perform complete case analysis or imputation prior to model fitting. Both approaches have serious drawbacks, potentially resulting in biased and unstable parameter estimates. Here we present `MGMM`, an `R` package for fitting GMMs in the presence of missing data. Using three case studies on real and simulated data sets, we demonstrate that, when the underlying distribution is near-to a GMM, `MGMM` is more effective at recovering the true cluster assignments than state of the art imputation followed by standard GMM. Moreover, `MGMM` provides an accurate assessment of cluster assignment uncertainty even when the generative distribution is not a GMM. This assessment may be used to identify unassignable observations. `MGMM` is available as an `R` package on CRAN: https://CRAN.R-project.org/package=MGMM.

## 1 Introduction

Gaussian mixture models (GMMs) provide a flexible approach to multivariate density estimation and probabilistic clustering (Murphy, 2012). Existing implementations of GMMs in the `R` programming language, such as `mclust` (Fraley and Raftery, 1999) and `mixtools` (Benaglia et al., 2009), require complete data. However, missing data are common in applications, potentially arising due to experimental reasons or when merging data sets with only partial variable overlap. Our development of this package was motivated by the problem of clustering summary statistics arising from genome-wide association studies (GWAS) of multiple correlated traits (Julienne et al., 2020a). Missing data arose because not every genetic variant was tested for association with every trait.

Although widely applied, standard approaches for addressing missing data prior to clustering, including complete case analysis and imputation, have serious drawbacks. By discarding information from observations that are only partially observed, complete case analysis makes inefficient use of the data. This leads to unstable estimates of model parameters and cluster assignments that would be susceptible to significant changes were the missingness pattern of the input data to change slightly. On the other hand, mean or median imputation introduces bias by making the incomplete observations appear less variable, and by shrinking the incomplete observations towards the complete data. This can result in inaccurate posterior membership probabilities that place excess weight on clusters with less missing data. Although methods have been described for estimating GMMs from incomplete data (Ghahramani and Jordan, 1994), there are no existing implementations in `R`.

To fill this gap, we present `MGMM`, an computationally efficient package for maximum likelihood estimation of GMMs in the presence of missing data. Our package is carefully implemented and documented for ease of use. Detailed examples of data generation and model fitting are provided in this article. In contrast to complete case analysis, our approach makes full use of the available data; and in contrast to clustering after imputation, our approach is unbiased for estimating the parameters of the generative GMM, accurately assesses the posterior membership probabilities, and correctly propagates estimation uncertainty.

Our implementation leverages the expectation conditional maximization (ECM) algorithm (Meng and Rubin, 1993), which accelerates estimation by breaking direct maximization of the EM objective function into a sequence of simpler conditional maximizations, each of which is available in closed form. While EM algorithms are regularly used for estimating GMMs, for example by both `mclust` and `mixtools`, those implementations only address missingness of the true cluster assignments, and not missingness of elements from the input vectors. In contrast, our ECM algorithm handles both missingness of the cluster assignments and of elements from the input data. We present a comprehensive benchmark, including three case studies, to demonstrate that when the underlying distribution is well-approximated by a GMM, `MGMM` is better able to recover the true cluster assignments than standard GMM applied after state of the art imputation, for example using multiple imputation by chained equations (MICE) (Buuren and Groothuis-Oudshoorn, 2010).

## 2 Models

This section describes the statistical model and details the ECM algorithm. The reader interested in the software may proceed to the next section.

### 2.1 Statistical model overview

Consider *n* independent vectors ***y***_*i*_ = vec(*Y*_*i*1_,⋯, *Y_id_*) in 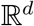, each arising from one of *k* distinct clusters. Let *Z_ij_* = 1 if the *i*th observation belongs to cluster *j*, and define the *k* × 1 indicator vector ***z***_*i*_ = vec(*Z*_*i*1_,⋯, *Z_ik_*). Conditional on membership to the *j*th cluster, ***y***_*i*_ follows a multivariate normal distribution, with cluster-specific mean ***μ***_*j*_ and covariance Σ_*j*_. Let *π_j_* denote the marginal probability of membership to the *j*th cluster. The observations can be viewed as arising from the following hierarchical model:

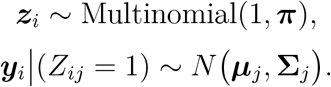

Marginalized over the latent cluster assignment vector ***z***_*i*_, each observation ***y***_*i*_ follows a *k* component Gaussian mixture model (GMM):

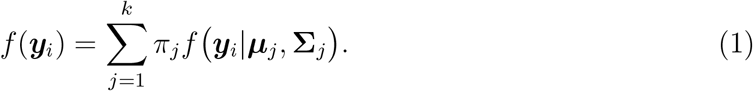

To perform estimation in the presence of missingness, we derive the EM objective function *Q*(***π, θ***|***π***^(*r*)^, ***θ***^(*r*)^), which is the expectation of the complete data log likelihood, given the observed data and current parameter estimates. The EM objective is optimized using a sequence of three conditional maximizations. Let 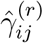 denote the *responsibility* of the *j*th cluster for the *i*th observation, which is the current conditional probability of membership to that cluster, given the observed data. In the first step, the cluster means are updated using the responsibility-weighted average of the working outcome vectors 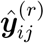. In the next step, the cluster covariances are updated using the responsibility-weighted average of the working residual outer products. In the final step, the cluster responsibilities and marginal membership probabilities are updated using the new means and covariances. This process iterates until the improvement in the EM objective drops below some tolerance *ϵ*.

### 2.2 Objective function

The log likelihood of the observed data is:

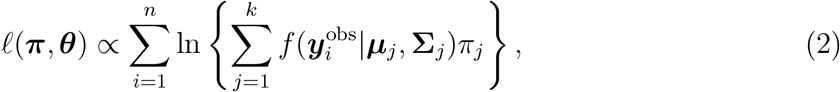

where 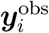 represents only the observed components of the complete-data vector ***y***_*i*_. However, due to the presence of a sum within the logarithm, there are no closed-form solutions for optimizing (2) in general. Here we develop the objective function optimized by the ECM algorithm, which is a lower bound on the observed data log likelihood (Hunter and Lange, 2004).

Define the *d* × *d residual outer product* matrix as:

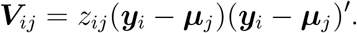

Let ***θ*** = (***μ***_1_, **Σ**_1_,⋯, ***μ***_*k*_, Σ_*k*_) collect the means and covariances of the component normal distributions. *Complete data* refers to the case where all elements of ***y***_*i*_ and all cluster assignments ***z***_*i*_ are observed. The complete data log likelihood is:

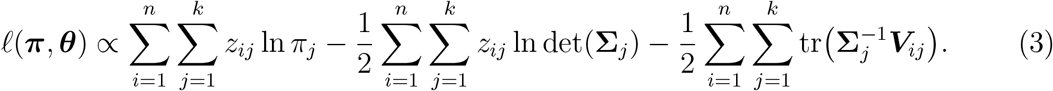

Let ***π***^(*r*)^ and ***θ***^(*r*)^ denote the current estimates of ***π*** and ***θ***, and let 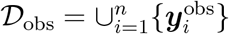 denote the observed data. The *ECM objective* is defined as the expectation of the complete data log likelihood (3) given the observed data and the current parameter states:

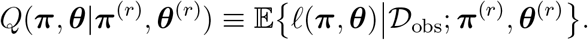

Define the *responsibility* of the *j*th cluster for the *i*th observation as the current conditional probability of membership given the observed data:

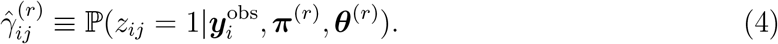

From Bayes’ theorem, the responsibility is expressible as:

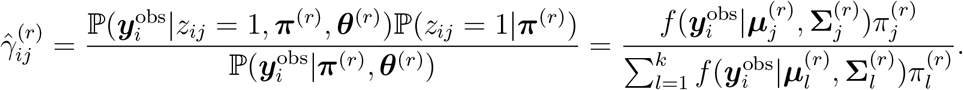

Here 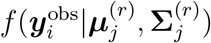 refers to the density of the observed elements 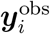 of ***y***_*i*_ given the current mean 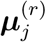 and covariance 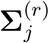 of the *j*th cluster.

Given membership to the *j*th cluster, the joint distribution of the observed 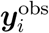 and missing 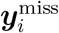 elements of ***y***_*i*_ is:

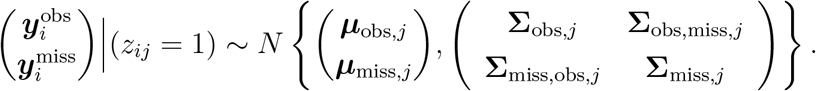

For the *i*th observation, define the *j*th *working response vector* as:

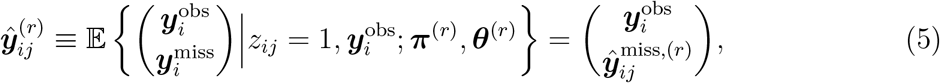

where 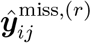 is the current conditional expectation of the missing elements of ***y***_*i*_ under the assumption that observation *i* originated from cluster *j*:

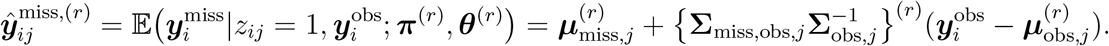

Finally, define the *working residual outer product* as:

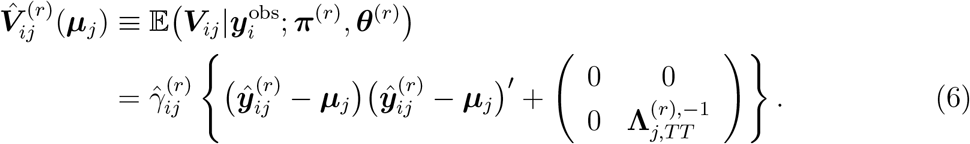

The working residual outer product is the current conditional expectation of the residual outer product given the observed data.

In terms of the responsibility and the working residual outer product, the ECM objective is:

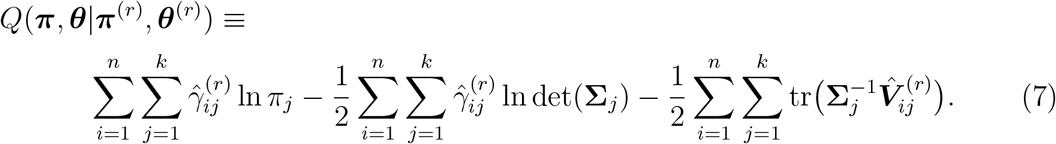

In contrast to the complete data log likelihood in (3), the ECM objective (7) is a function of the observed data only.

### 2.3 Optimization

ECM differs from classic EM in that the M-step is partitioned into a sequence of conditional maximizations (Meng and Rubin, 1993). In the present case, estimation of the means (***μ***_*j*_), followed by the covariances (Σ_*j*_), and finally the cluster membership probabilities ***π***. The advantage of ECM is that each conditional maximization is available in closed form.

The optimization procedure is outlined in Algorithm 1. Initial estimates of ***π*** and ***θ*** are required. These may be obtained, for example, by performing an initial *k*-means clustering on complete observations (Bishop, 2006), then calculating the within-cluster means and covariances. Update equations were derived by differentiating the ECM objective in (7) and solving the resulting score equations. The update for ***μ***_*j*_ is:

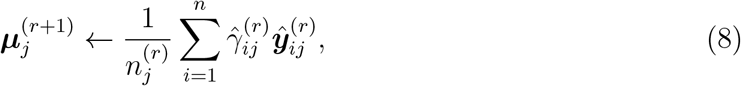

where 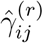 is the current responsibility (4) of the *j*th cluster for the *i*th observation,

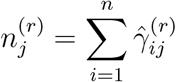

is the total responsibility of the *j*th cluster, and 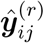 is the current working response vector (5). Note that (8) takes the form of a responsibility-weighted average of the working response vectors. Similarly, the update for **Σ**_*j*_ is:

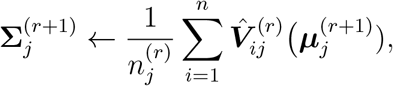

which is a responsibility-weighted average of the working residual outer products (6).

Given the updated means 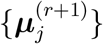 and covariates 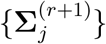, the new responsibilities are:

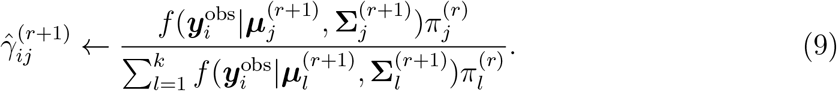

Finally, the update for *π_j_* is:

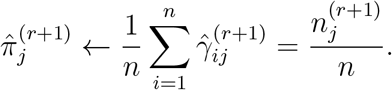

After all parameters have been updated, the current value of the ECM objective *Q*^(*r*+1)^ = *Q*(***π***^(*r*+1)^, ***θ***^(*r*+1)^|***π***^(*r*+1)^, ***θ***^(*r*+1)^) is calculated from (7), and the algorithm continues until the increment *Q*^(*r*+1)^ – *Q*^(*r*)^ in the objective falls below a pre-specified tolerance *ϵ*.

**Algorithm 1.**
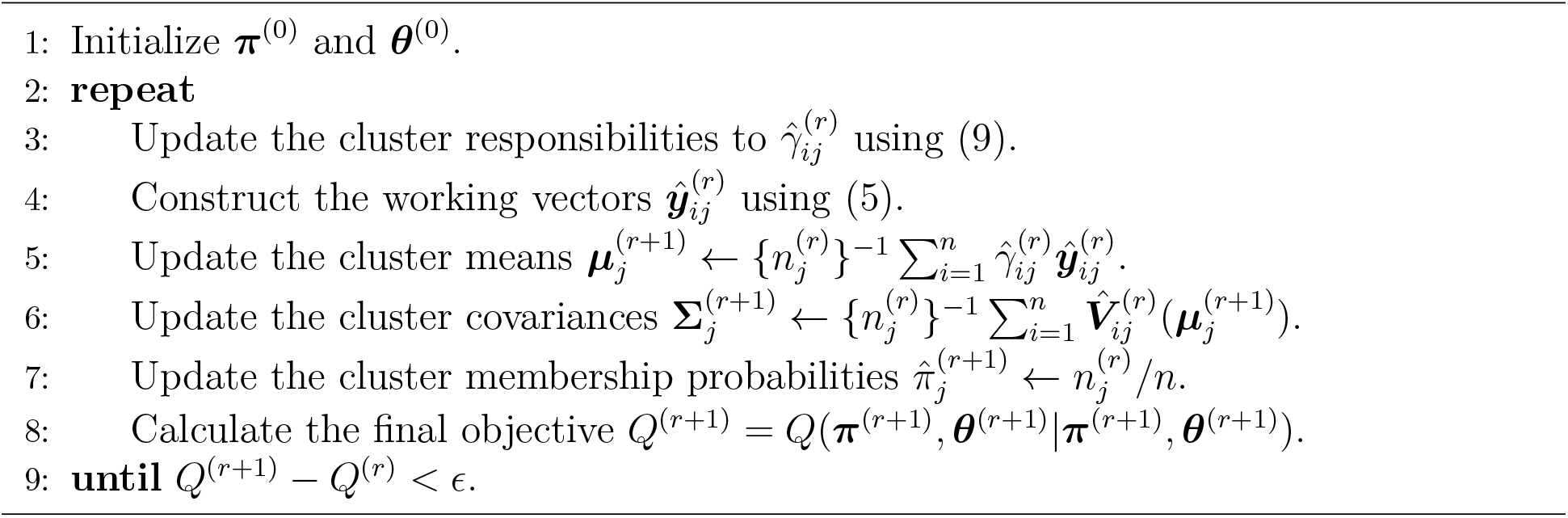
ECM for GMMs

### 2.4 Imputation

Having fit the GMM in (1) via maximum likelihood, the missing values 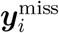 of each observation ***y***_*i*_ may subsequently be imputed to their posterior expectations. Note that, in contrast to the comparators described below, this imputation is occurring *after* estimation, and has no effect on the final maximum likelihood estimates 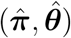. The posterior expectation of the missing data is:

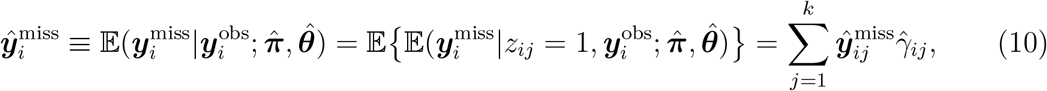

where 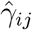 is the final responsibility of cluster *j* for observation *i*, and:

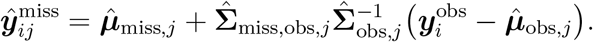

### 2.5 Missingness

Standard approaches to addressing missing data, including complete case analysis and naive mean or median imputation, often tacitly assume that the data are missing completely at random (MCAR). Under MCAR, whether an element of data is missing is completely independent of its value. However, unbiased estimation via the maximum likelihood-based approach presented here only requires that the missingness occurs at random (MAR), which is a weaker and more plausible assumption. Under MAR, whether an element of data is missing is independent of its value conditional on those elements of the data that are observed (Little and Rubin, 2002).

To elaborate on this assumption, let *R_il_* = 1 if the *l*th component of the *i*th observation (that is, *Y_il_*) is observed, and define the response indicator vector ***r***_*i*_ = vec(*R*_*i*1_,⋯, *R_ip_*). For the *i*th subject, let 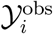 denote the indices of the elements of ***y***_*i*_ that are observed, and 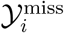 the indices of the elements that are missing. Partition ***y***_*i*_ as 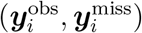, where 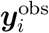 contains the observed elements of ***y***_*i*_, and 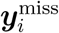 contains the missing elements. The MAR assumption requires that, for each observation *i* and all missing elements 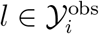, the response indicator *R_il_* is conditionally independent of *Y_il_* given that observation’s observed data 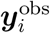. For example, MAR holds under the generalized linear model 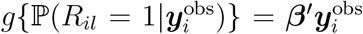, for some regression coefficient ***β***. MAR fails if the missing occurs not at random (MNAR), for example if 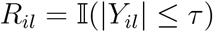 for some threshold *τ*, or if 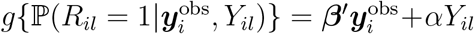 for *α* ≠ 0. That is, the data are MNAR if 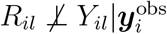.

## 3 R Package Functionalities and Usage Examples

### 3.1 Data Simulation

#### Description

The function `rGMM` simulates observations from a Gaussian mixture model (1). The number of observations is specified by `n`, and the dimension of each observation by `d`. The number of clusters is set using `k`, which defaults to one. The marginal probabilities of cluster membership are provided as a numeric vector `pi`, which should contain `k` elements. If *k* > 0 is provided but `pi` is omitted, the clusters are assumed equiprobable. The proportion of elements in the *n*×*d* data matrix that are missing is specified by `miss`, which defaults to zero. Missingness is introduced completely at random. Note that when miss > 0 it is possible for all elements of an observation to be missing. The cluster `means` are provided either as a numeric prototype vector, or a list of such vectors. If a single prototype is provided, that vector is used as the mean of all clusters. By default, the zero vector is adopted as the prototype. The cluster covariances (`covs`) are provided as a numeric prototype matrix, or a list of such matrices. If a single prototype is provided, that matrix is used as the covariance for all clusters. By default, the identity matrix is adopted as the prototype.

##### **Examples** Single Component without Missingness

In this example, `n = 1e3` observations are simulated from a single `k = 1` bivariate normal distribution `d = 2` without missingness (`miss = 0`, by default). The mean is *μ* = (2, 2), and the covariance is an exchangeable correlation structure with off-diagonal *ρ* = 0.5.

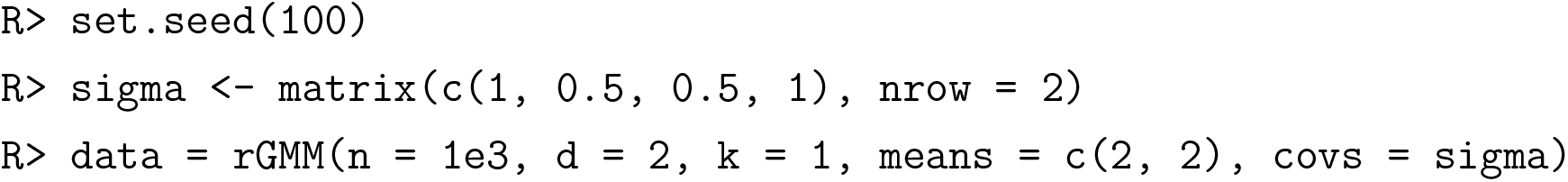

##### Single Component with Missingness

Here, `n = 1e3` observations are simulated from a single `k = 1` trivariate normal distribution `d = 3` with 20% missingness `miss = 0.2`. The mean defaults to the zero vector, and the covariance to the identity matrix.

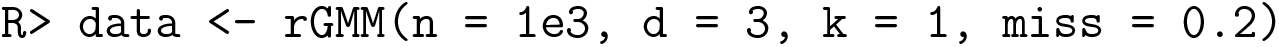

##### Two Components without Missingness

In this example, `n = 1e3` observations are simulated from a two-component `k = 2` trivariate normal distribution `d = 3` without missingness. The mean vectors are *μ*_1_ = (−2, −2, −2) and *μ*_2_ = (2, 2, 2). The covariance matrices are both exchangeable with off-diagonal *ρ* = 0.5. Since `pi` is omitted, the cluster are equiprobable, i.e. ***π*** = (0.5, 0.5).

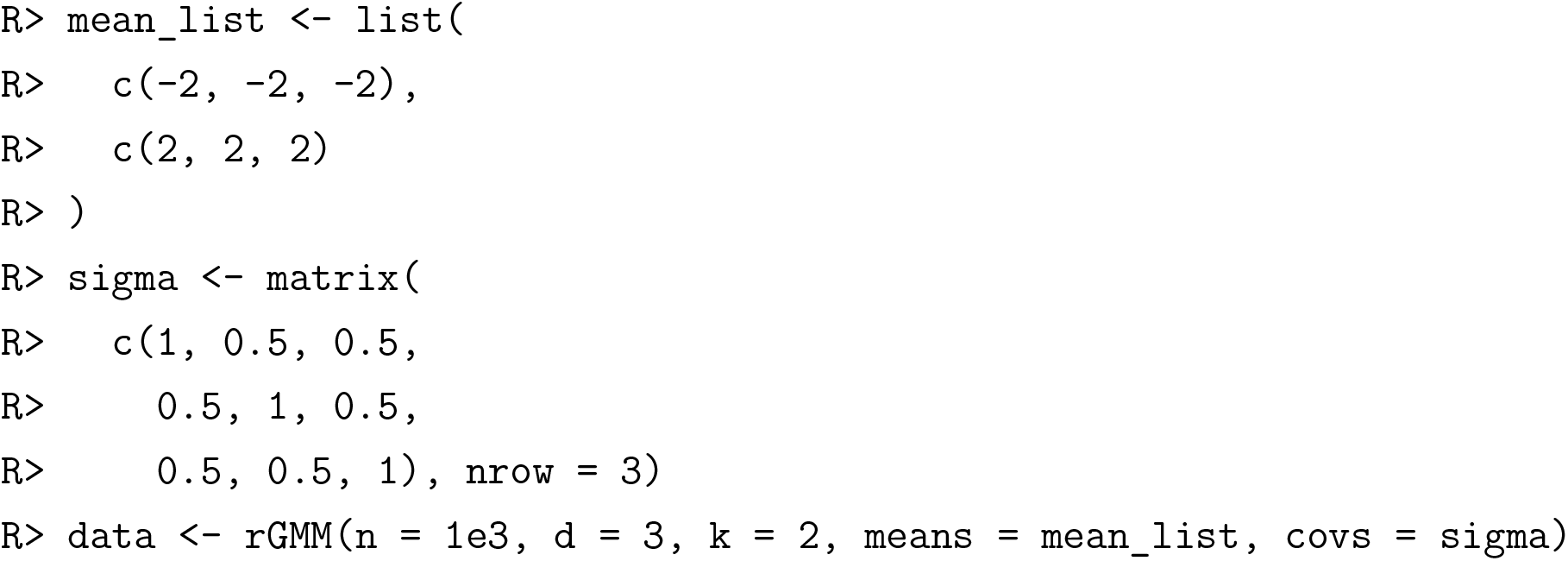

##### Four Components with Missingness

Here, `n = 1e3` observations are simulated from a four-component `k = 4` bivariate normal distribution `d = 2` with 10% missingness `miss = 0.1`. The mean vectors are *μ*_1_ = (−2, −2), *μ*_2_ = (−2, 2), *μ*_3_ = (2, −2) and *μ*_4_ = (2, 2). The covariance matrices are all 0.5 * *I*. The cluster proportions are (35%, 15%, 15%, 35%) for (*π*_1_, *π*_2_, *π*_3_, *π*_4_), respectively.

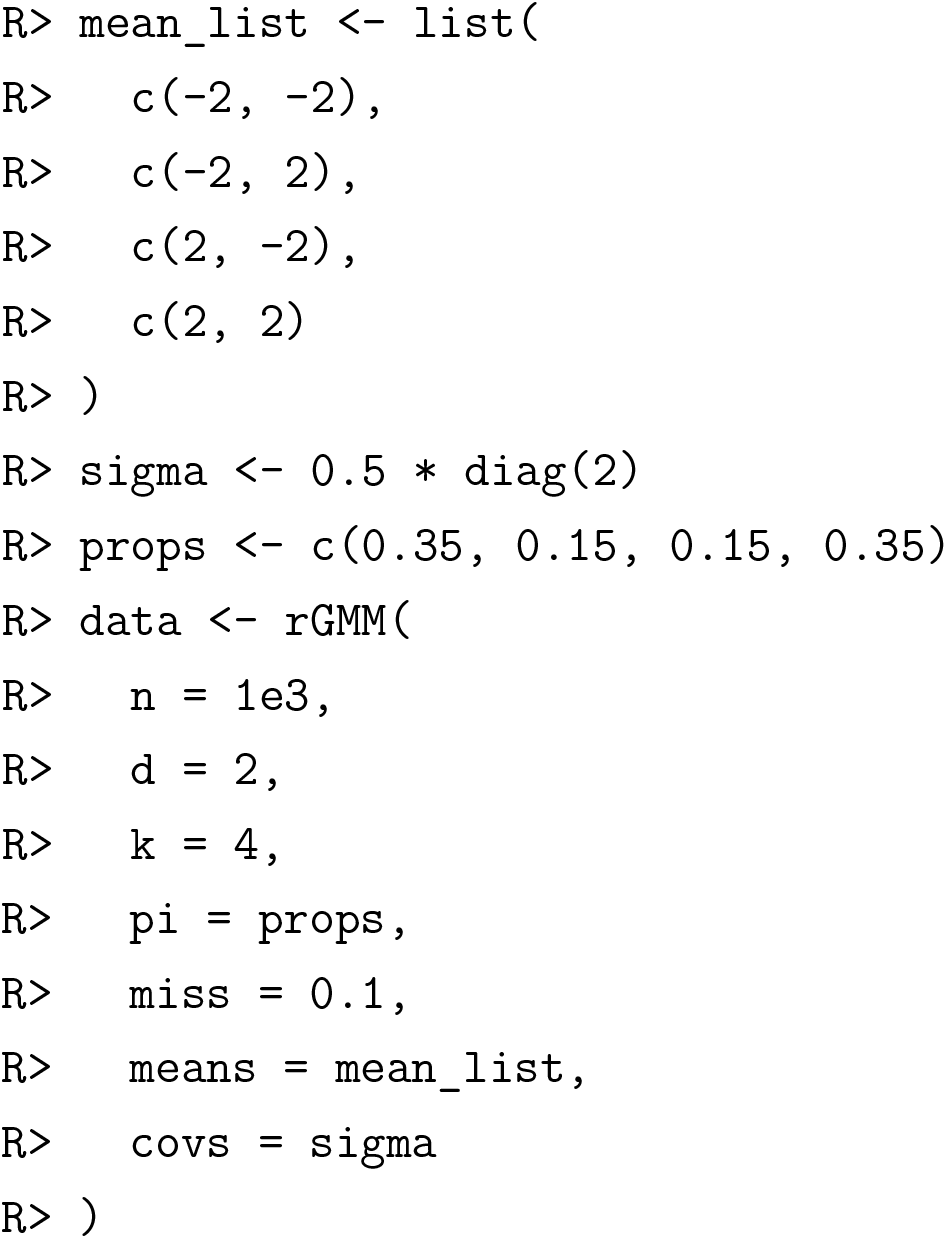

### 3.2 Parameter Estimation

#### Description

The function `fit.GMM` estimates the parameters of the GMM. The data are expected as a numeric matrix `data`, with independent observations as rows. The number of mixture components is specified using `k`, which defaults to one. Initial values for the mean vectors, covariance matrices, and cluster proportions are provided using `init_means, init_covs`, and `init_props`, respectively. If the `data` contain complete observations, i.e. observations with no missing elements, `fit.GMM` will attempt to initialize the model parameters (*μ*, Σ, *π*) via K-means. However, if the data contain no complete observations, initial values are required for each of `init_means, init_covs`, and `init_props`. Supplying initial values may also accelerate estimation when there are relatively few complete observations. Initial means and covariances should be supplied as lists with k components, even if there is a single component (*k* = 1), or all *k* > 1 components are initialized at the same value.

The arguments `maxit, eps`, and `report` control the fitting procedure. `maxit` sets the maximum number of ECM iterations to attempt. The default is 10^2^. `eps` sets the minimum acceptable improvement in the ECM objective function. The default is 10^-6^. If `report = TRUE`, then fitting progress is displayed.

#### Examples

##### Single Component without Missingness

In this example, `n = 1e3` observations are simulated with a single bivariate normal distribution without missingness. Since the model contains only a single component, the output is a list containing the estimated mean and covariance. In the case of a single component without missingness, the maximum likelihood estimates are available in closed form.

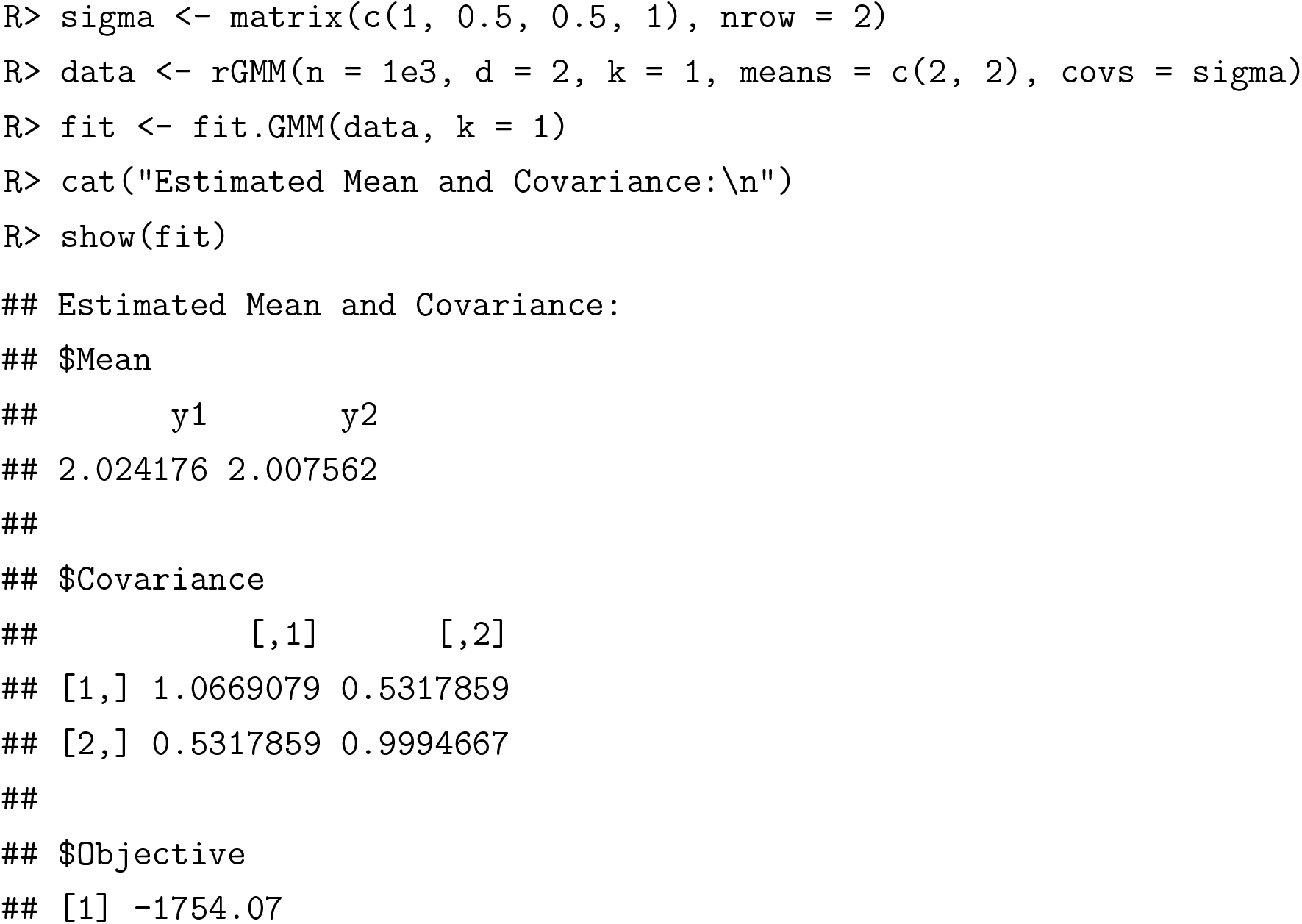

##### Single Component with Missingness

Here, `n = 1e3` observations are simulated from a single trivariate normal distribution with 20% missingness. Since the model contains only a single component, the output is again a list. However, in the case of missingness, the ECM algorithm is used for estimation, and a completed version of the input data is returned, with missing values imputed to their posterior expectations. The true mean is the zero vector, and the true covariance is identity. For `fit1` below, the initial mean and covariance are estimated internally using complete observations. For `fit2` below, the mean and covariance are initialized at the truth.

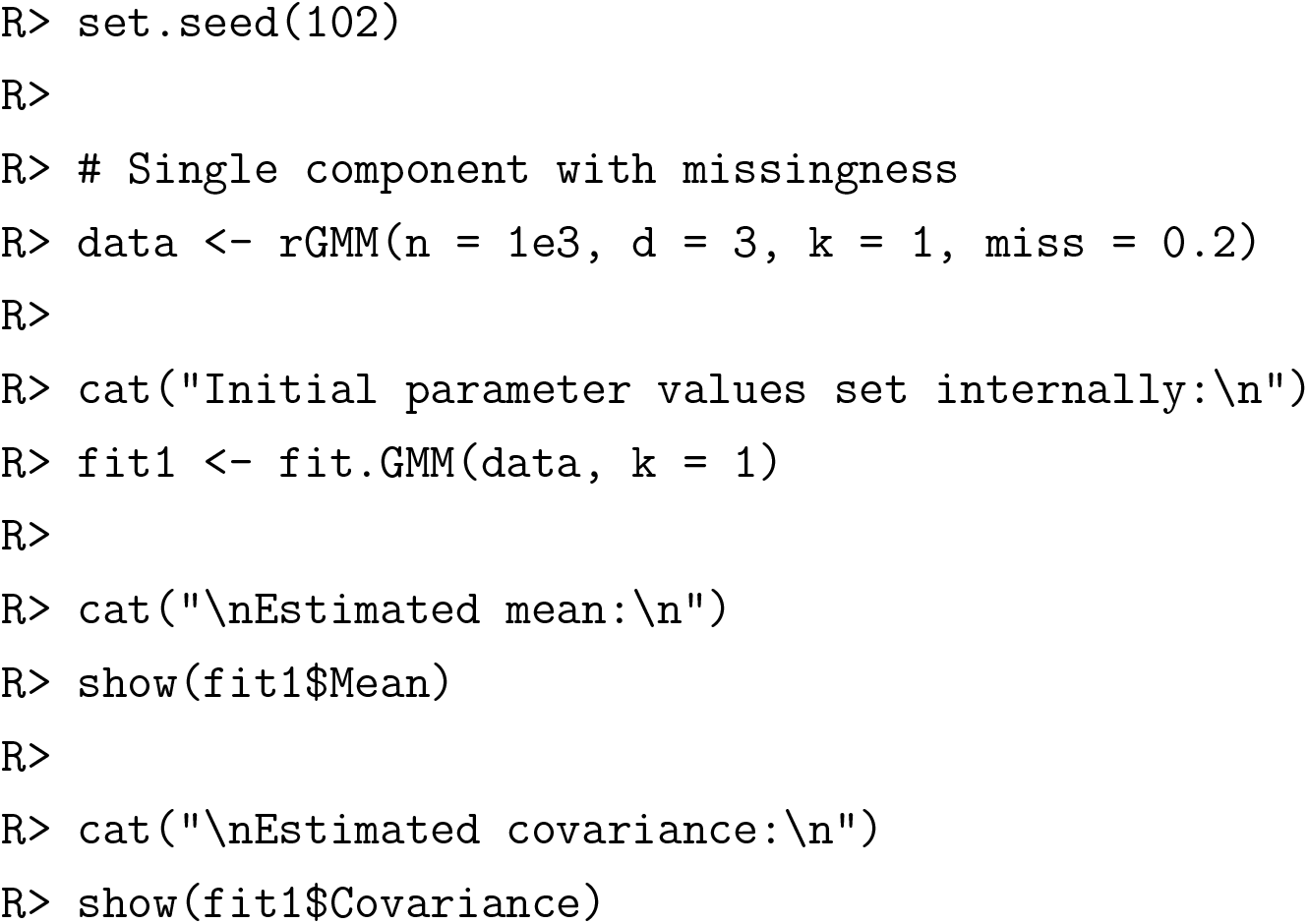
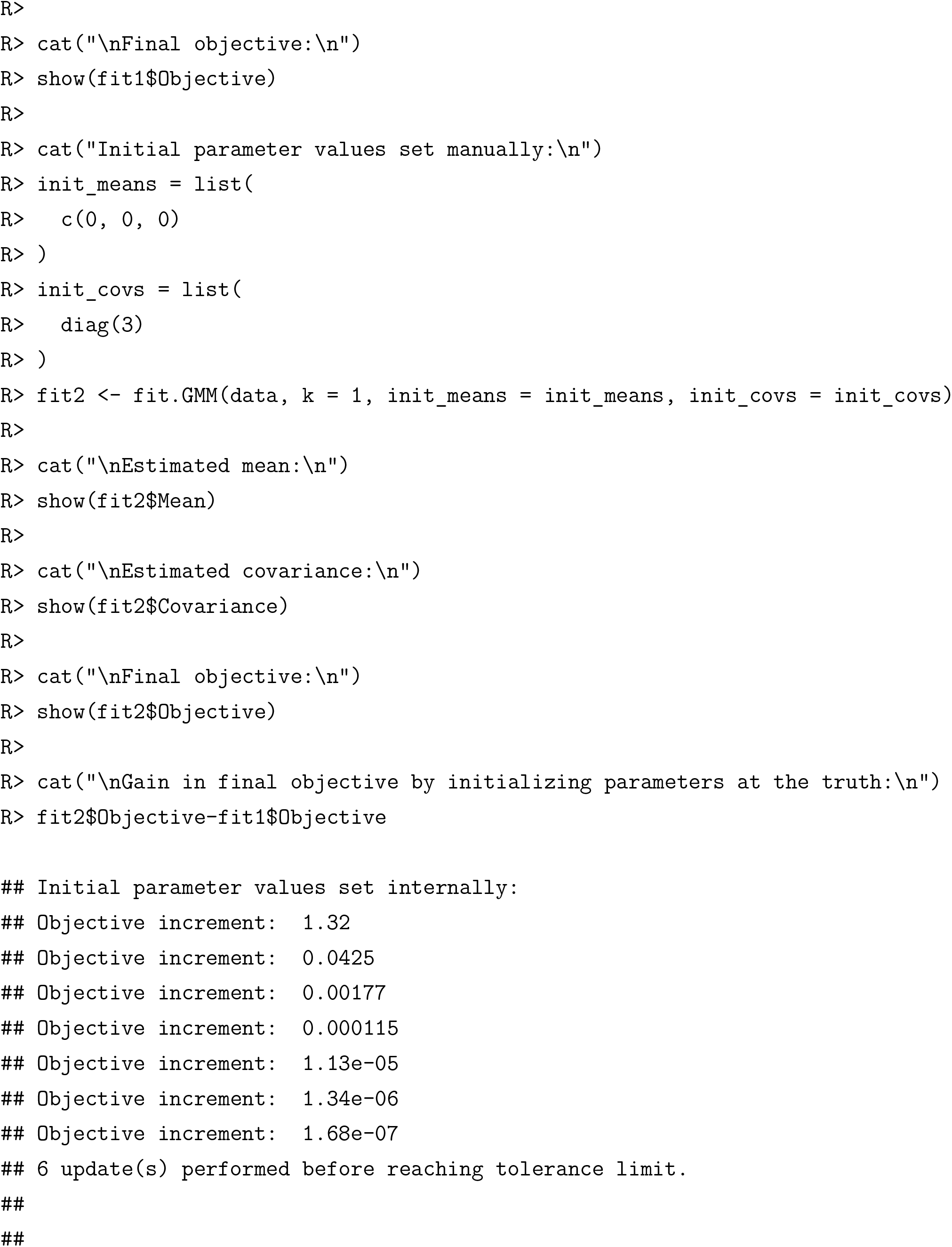
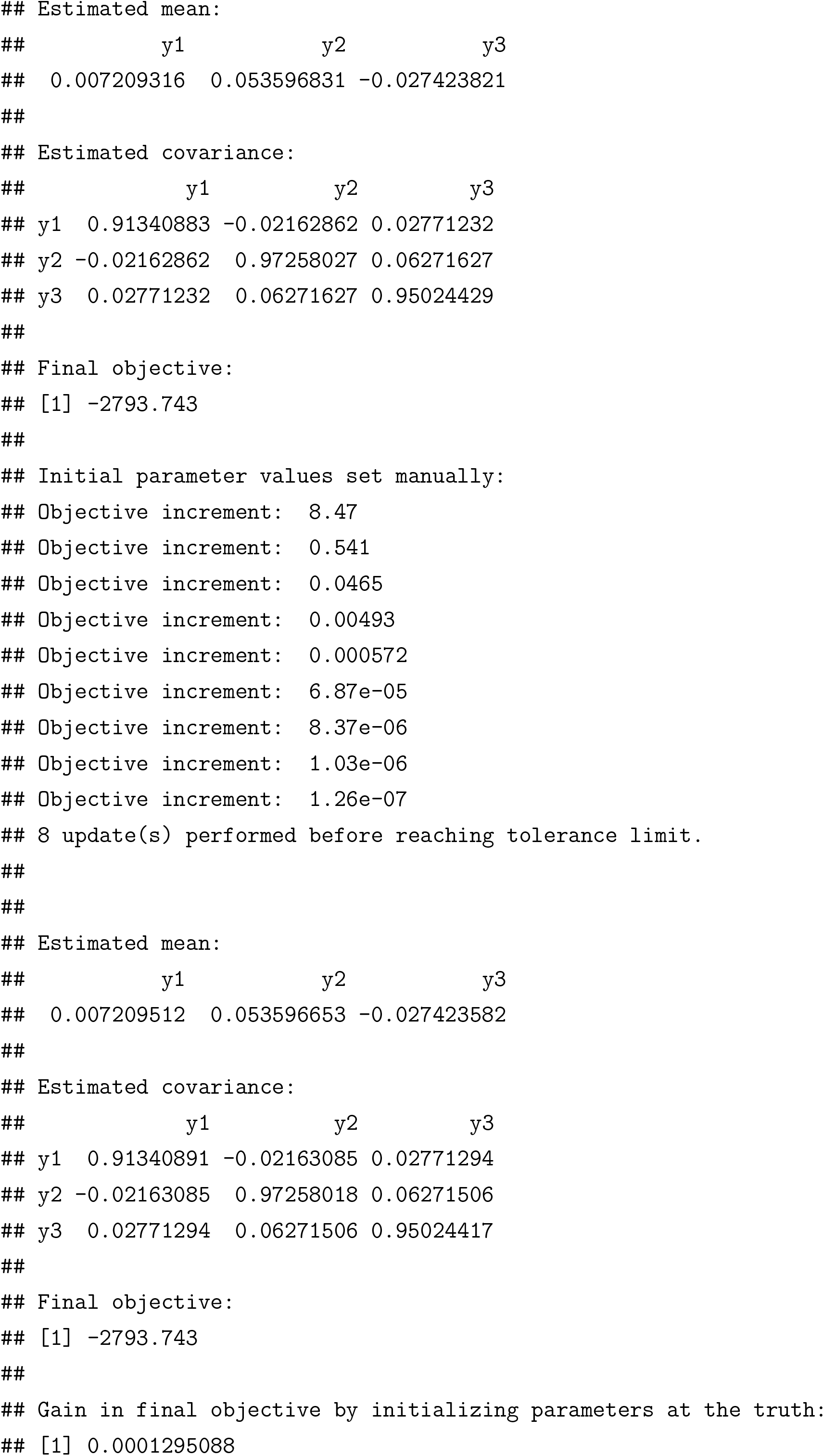

##### Two Components without Missingness

In this example, `n = 1e3` observations are simulated from a two-component, trivariate normal distribution without missingness. Since the model has multiple components, the output is an object of class `mix`. The `show` method displays only the estimated cluster proportions and the final objective. The slots of the output contain the following:

- @Means and @Covariances: lists of the estimated cluster means and covariances.
- @Density: the cluster densities evaluated at the observations, which may be useful for identifying outliers, as an outliers will tend to have little support from any cluster.
- @Responsibilities: the posterior probabilities of membership to each cluster.
- @Assignments: the maximum *a posteriori* cluster assignments, and the assignment entropy. A lower assignment entropy indicates higher confidence cluster assignments.
- @Completed: a completed version of the input data is returned, with missing values imputed to their posterior expectations (10). Note that this imputation is occurring after the maximum likelihood estimation.

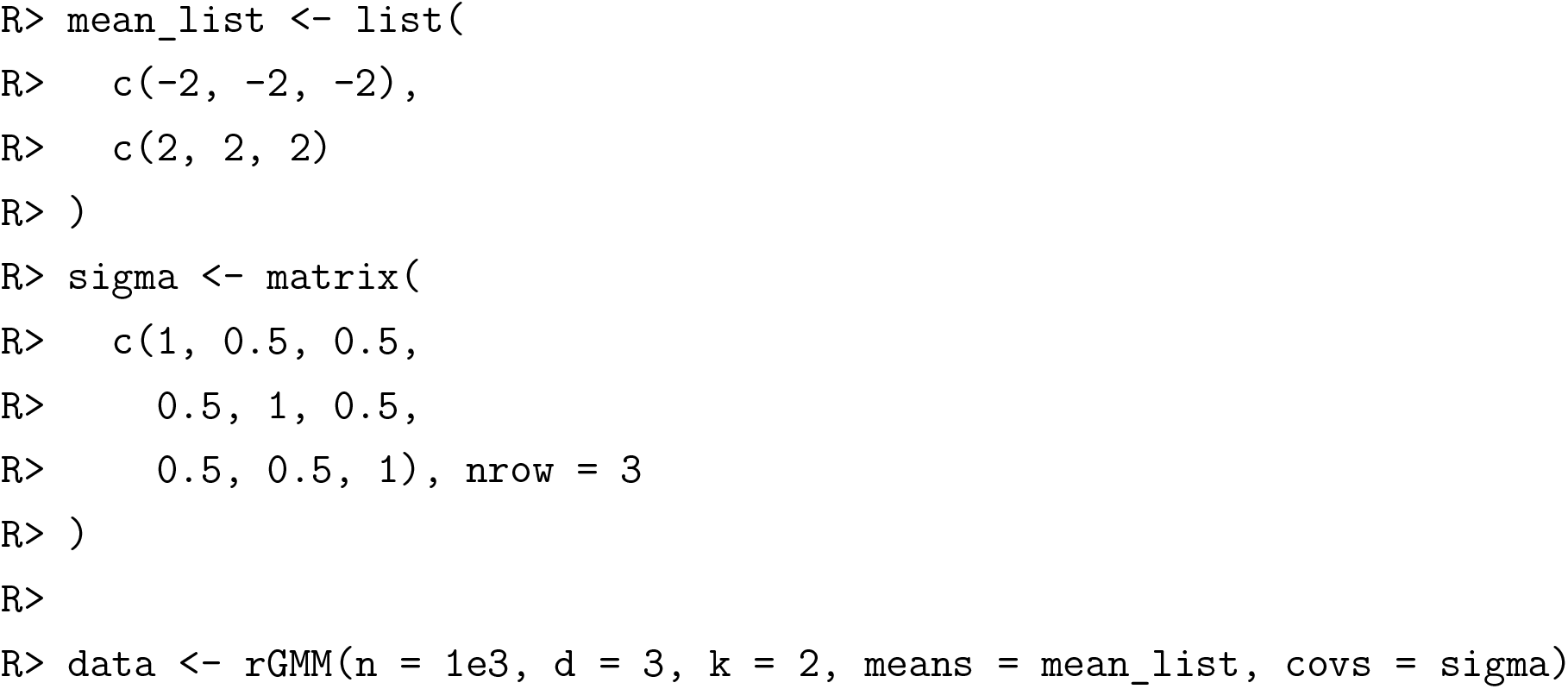
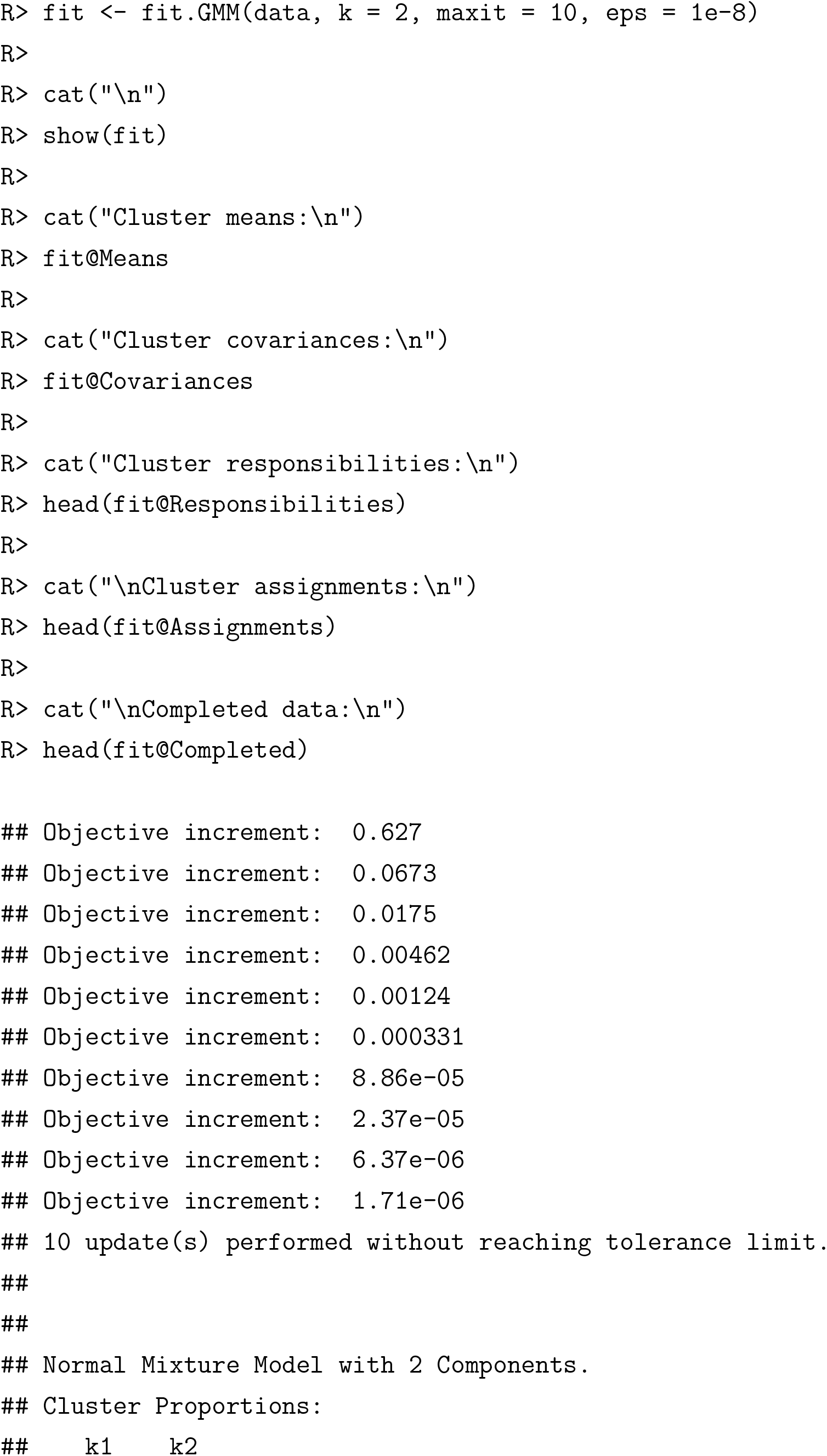
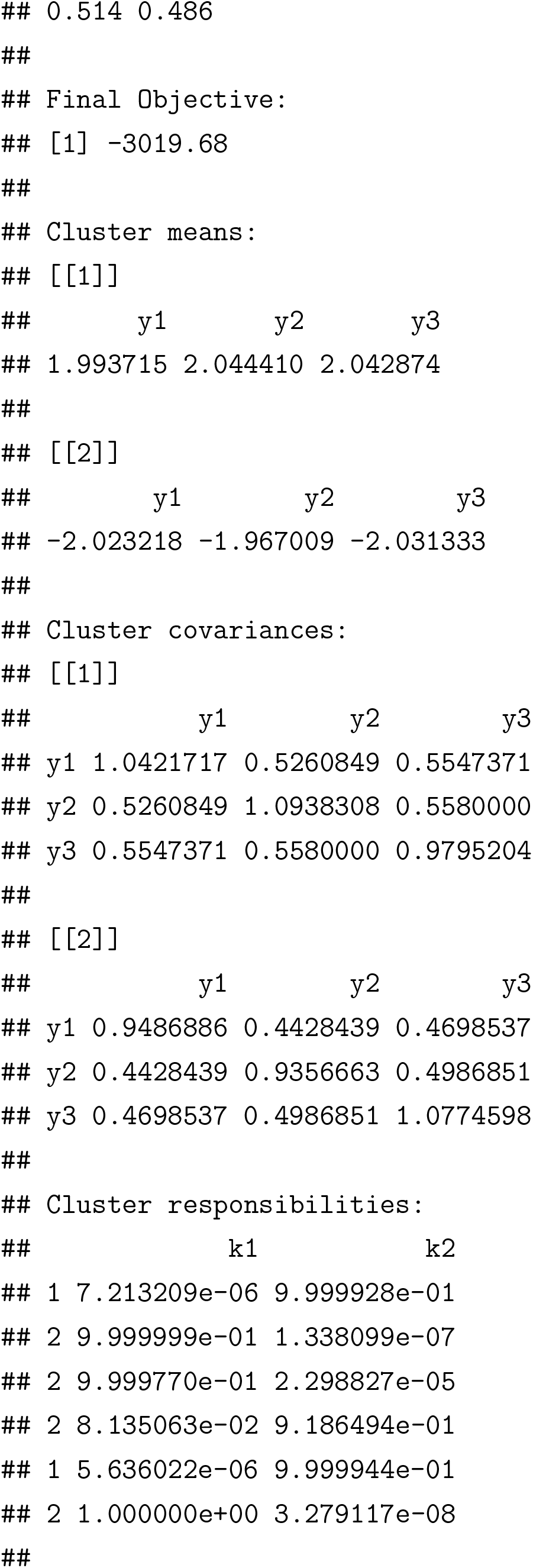
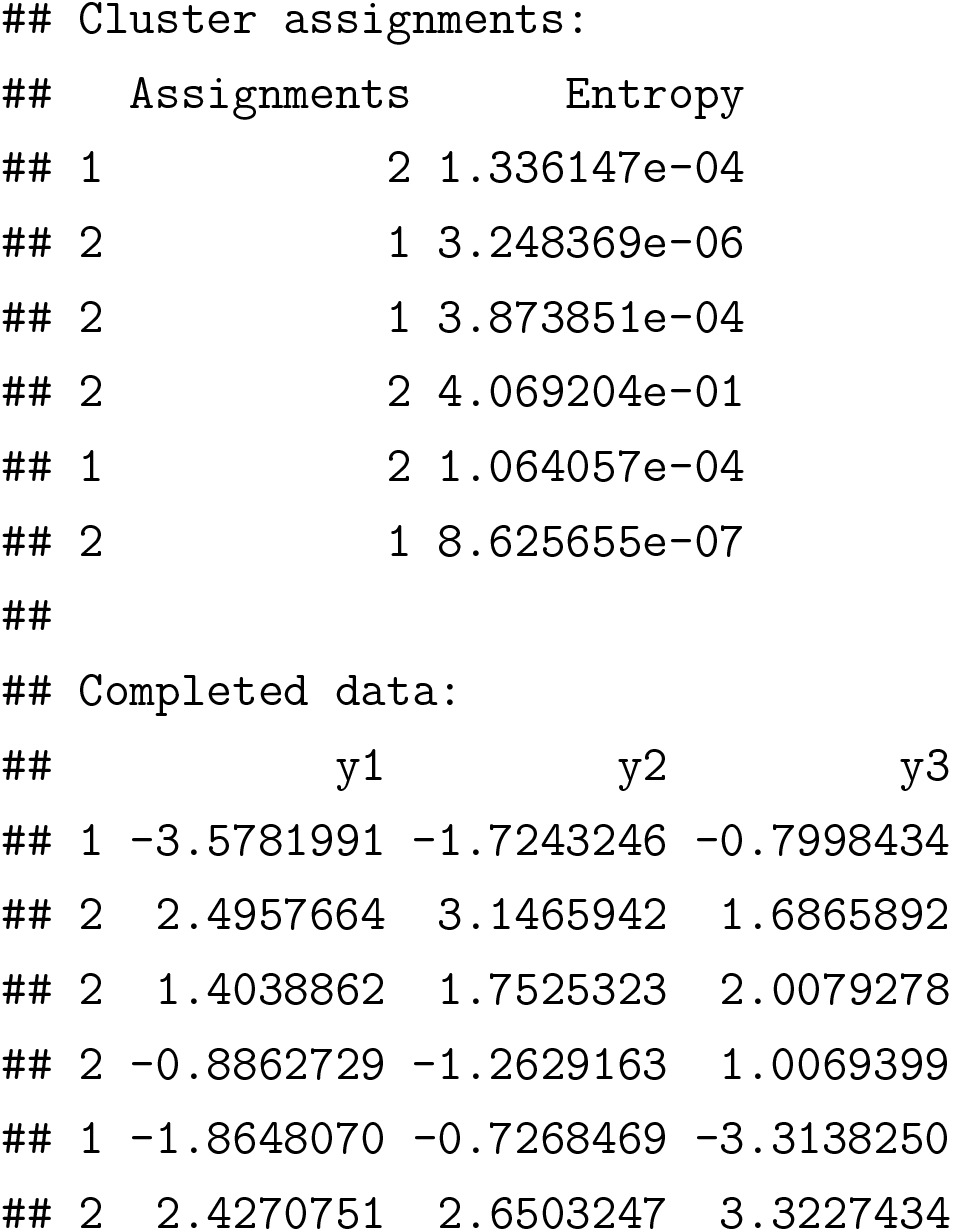

##### Four Components with Missingness

Here, `n = 1e3` observations are simulated from a four-component bivariate normal distribution with 10% missingness. Since the model has multiple components, the output is an object of class `mix`.

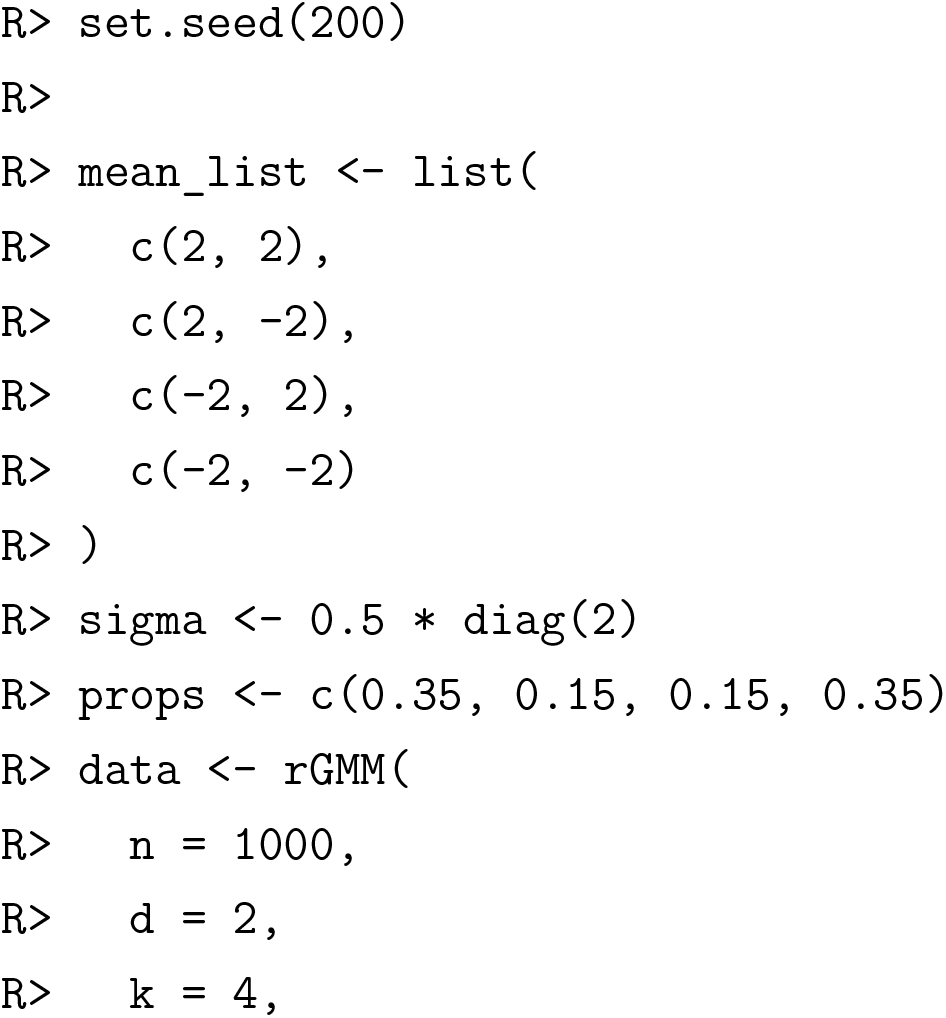
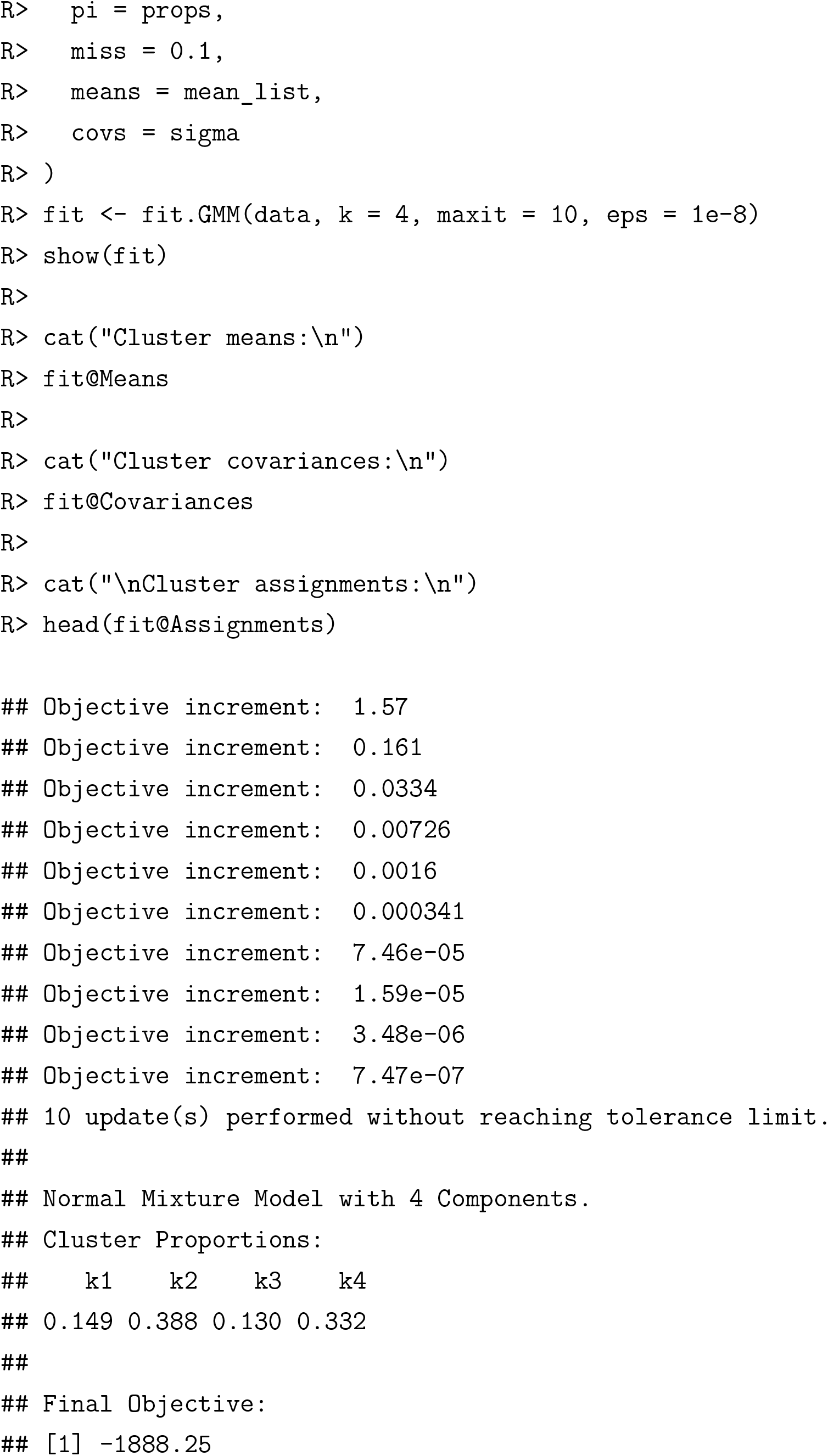
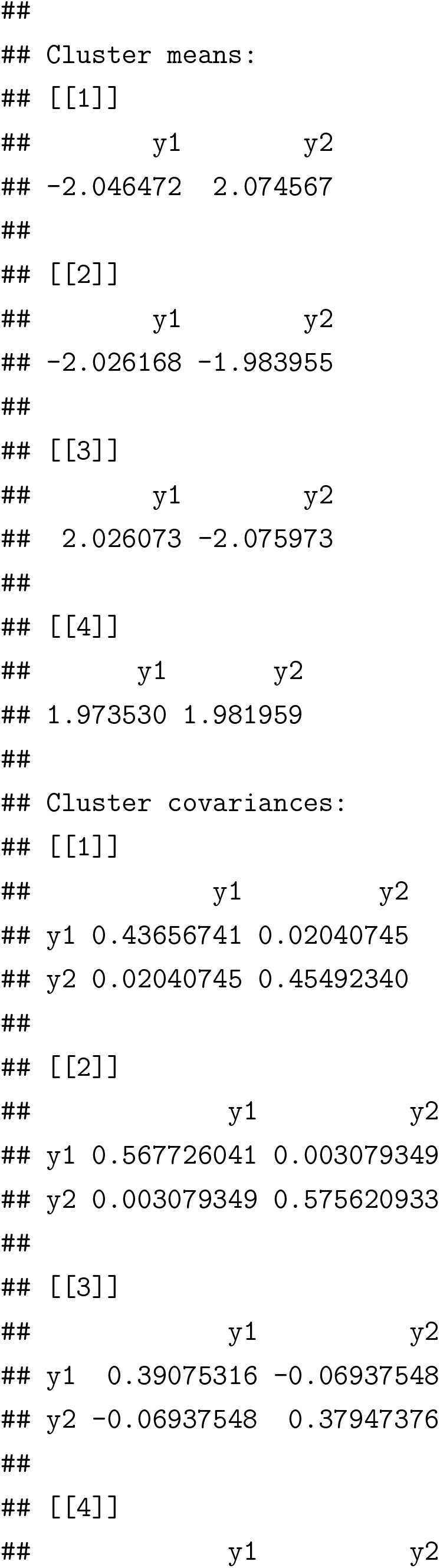
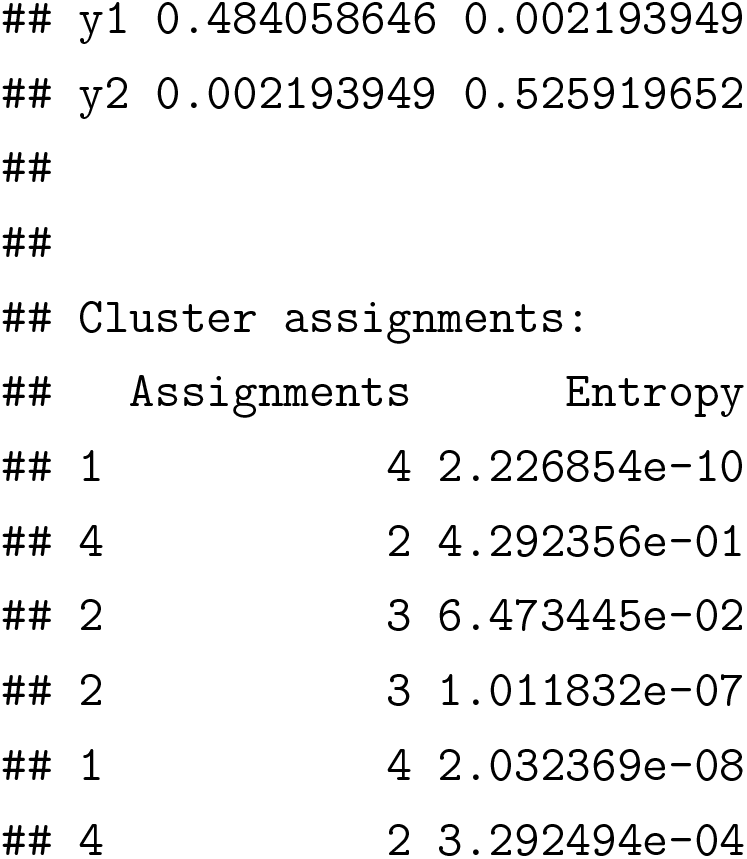

### 3.3 Cluster Number Selection

#### Clustering Quality

The function `ClustQual` provides several metrics for internally assessing the quality of cluster assignments from a fitted GMM. The input is an object of class `mix`. The output is a list containing these metrics: `BIC, CHI, DBI`, and `SIL`.

- `BIC` is the Bayesian information criterion, which is a penalized version of the negative log likelihood. A lower value indicates better clustering quality.
- `CHI` is the Calinski-Harabaz index, a ratio of the between-cluster to within-cluster variation. A higher value indicates better clustering quality.
- `DBI` is the Davies-Bouldin index, an average of similarities across clusters. A lower value indicates better clustering quality.
- `SIL` is the average silhouette width, a measure of how well an observation matches its assigned cluster. A higher value indicates better clustering quality.

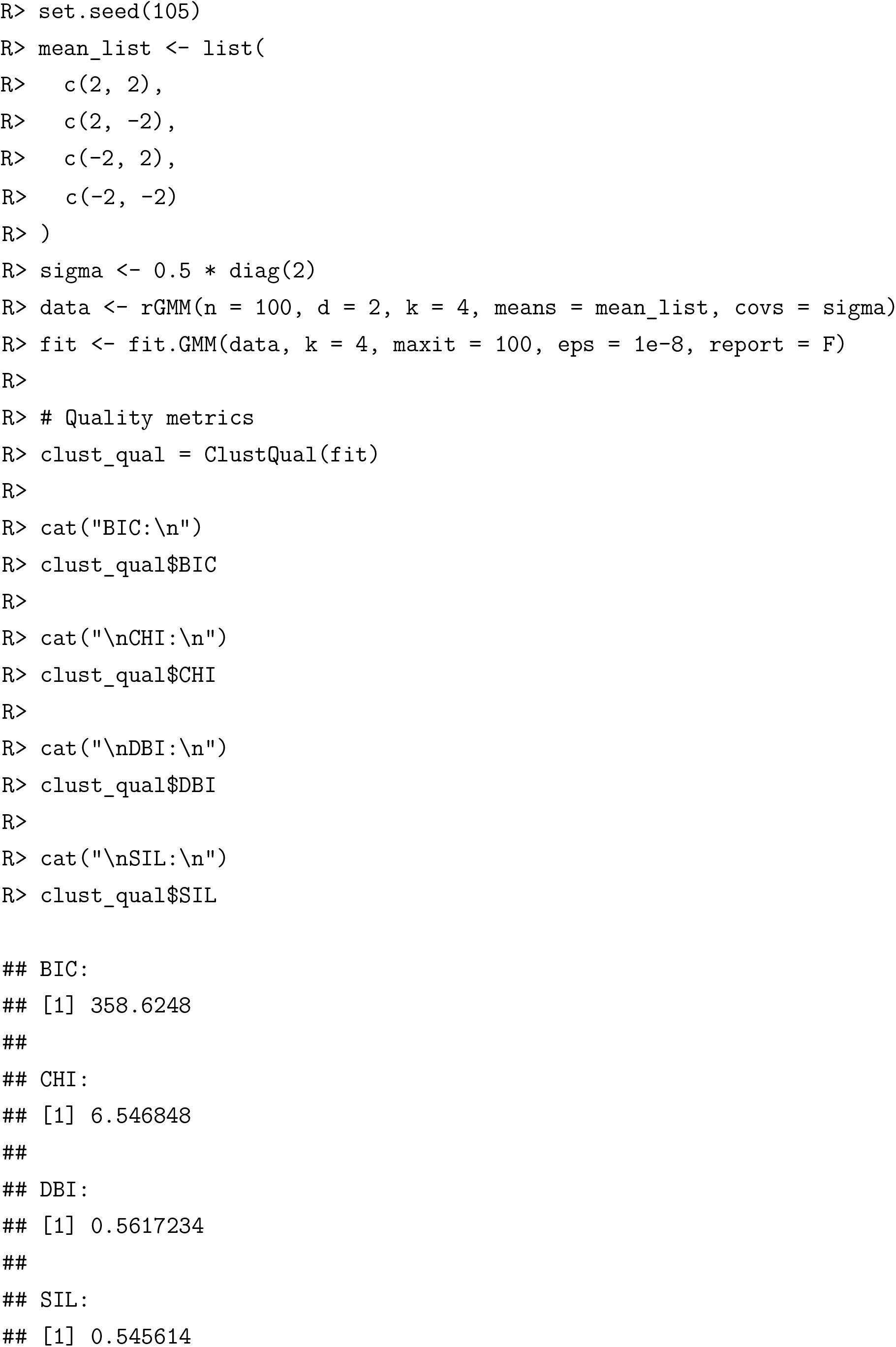

#### Choosing the Number of Clusters

In applications, the number of clusters `k` is often unknown. The function `ChooseK` is designed to provide guidance on the number of clusters present in the data. The inputs include the data matrix `data`, the minimum number of clusters to assess `k0`, the maximum number of clusters to assess `k1`, and the number of bootstrap replicates at each cluster number `boot`. For each cluster number `k` from `k0` to `k1`, boot bootstrapped data sets are generated. A GMM with `k` components is fit, and the quality metrics are calculated. The bootstrap replicates are summarized by their mean and standard error (SE). For each quality metric, the cluster number `k_opt` that provided the optimal quality, and the smallest cluster number whose quality was within 1 SE of the optimum `k_1se`, are reported. The output is a list containing two data frames:

- `Choices`, summarizing `k_opt` and `k_1se` for each cluster quality metric.
- `Results`, reporting the mean and standard error of each cluster quality metric at each cluster size evaluated.

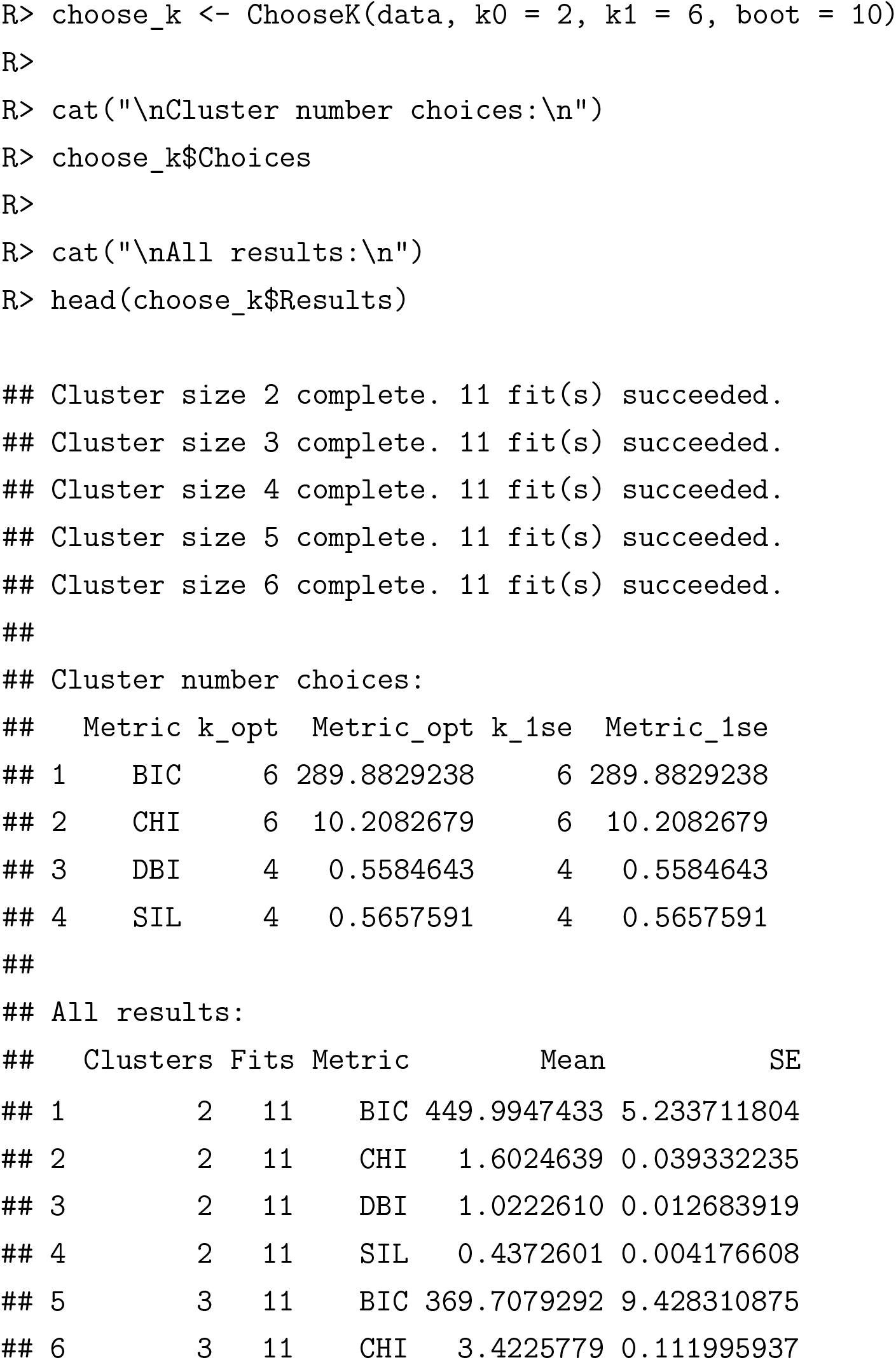

## 4 MGMM vs. State of the Art Imputation Methods

We designed a benchmarking procedure to compare the performance of `MGMM` against imputation followed by standard GMM. We defined clustering performance as the capacity of the algorithm to recover the true cluster assignments when applied to example data sets. We assessed the quality of the clustering by calculating the adjusted Rand index (ARI) between the recovered and true class assignments. The imputation methods included in the benchmark were: naive mean and median imputation, k-nearest neighbors (k-NN) imputation (Kowarik and Templ, 2016), multiple imputation by chained equations (MICE) (Buuren and Groothuis-Oudshoorn, 2010), and random forest imputation (Stekhoven and Bühlmann, 2011). We applied the benchmarking procedure to four case studies: a simulated four component mixture of bivariate Gaussians, a cancer patient RNA-seq data set, simulated genome-wide association studies (GWAS) summary statistics, and summary statistics from GWAS for cardiovascular disease risk factors. (Julienne et al., 2020b).

### 4.1 Benchmarking Method

#### 4.1.1 Missingness

For *n* observations on *d* dimensional data, a fraction of missing values *m* was introduced completely at random by setting ⌉(*m* × *n* × *d*)⌈ elements of the data set to `NA`.

#### 4.1.2 Evaluation Metric

The quality of clustering was evaluated using the ARI Rand (1971); Hubert and Arabie (1985). Briefly, the Rand index (RI) is a measure of similarity that assesses the agreement between two partitions of a collection of *n* objects. All possible pairs of objects are examined, and the proportion of pairs that are either 1. in the same cluster or 2. in different clusters according to both partitions is calculated. ARI is a variation of the RI that is adjusted for chance, and is permutation invariant. A value near zero suggests the agreement between the two partitions is no better than expected by chance, and a value of one occurs when the two partitions are identical. We define the quality of clustering as the ARI between the reference or true clustering, established in the data set description, and the clustering performed in the presence of missingness.

#### 4.1.3 Benchmarking Procedure

We designed the benchmarking procedure outlined in Figure 1 and described in Algorithm 2 to compare the performance of `MGMM` with imputation followed by standard GMM.

**Figure 1:**
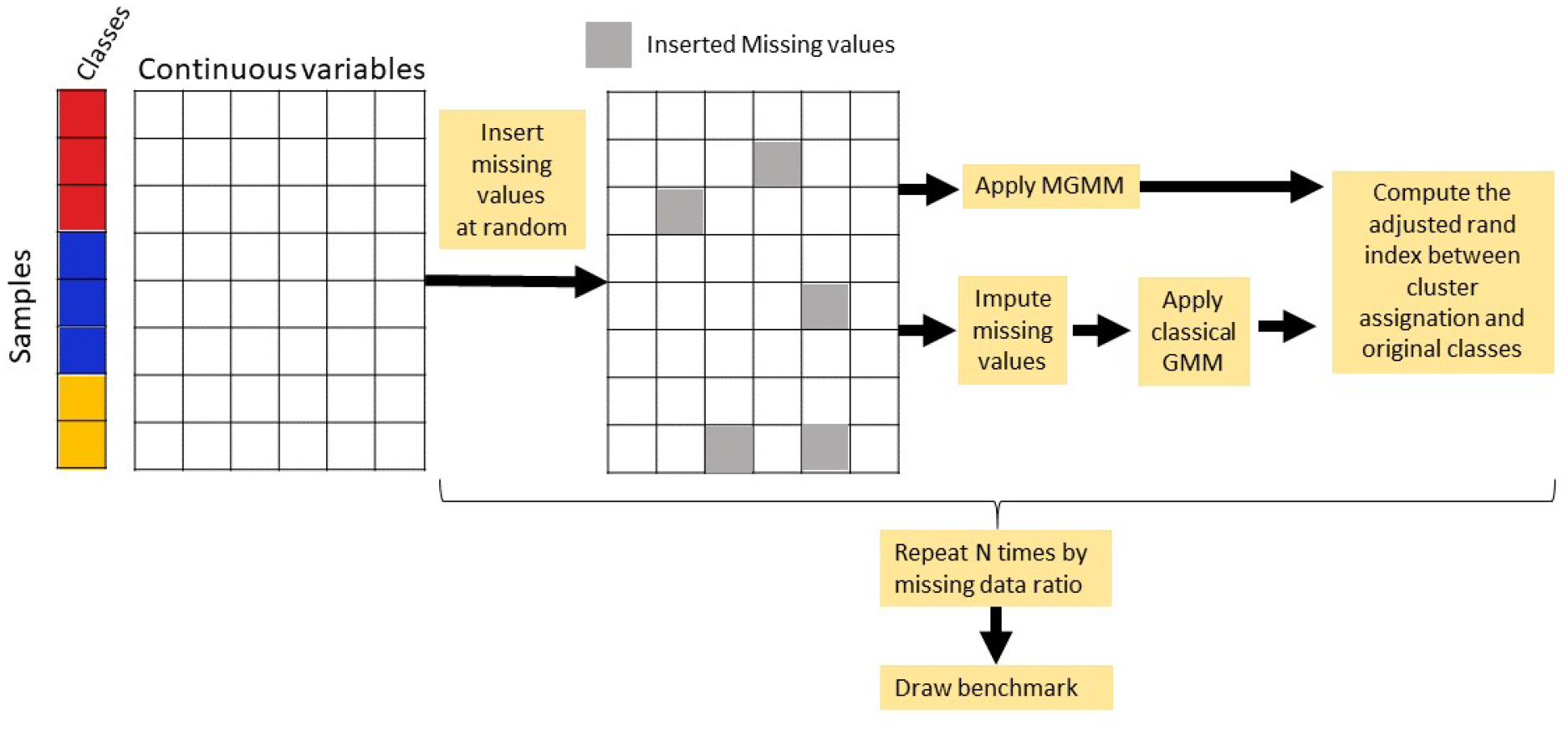
Benchmark procedure schematics

**Algorithm 2.**
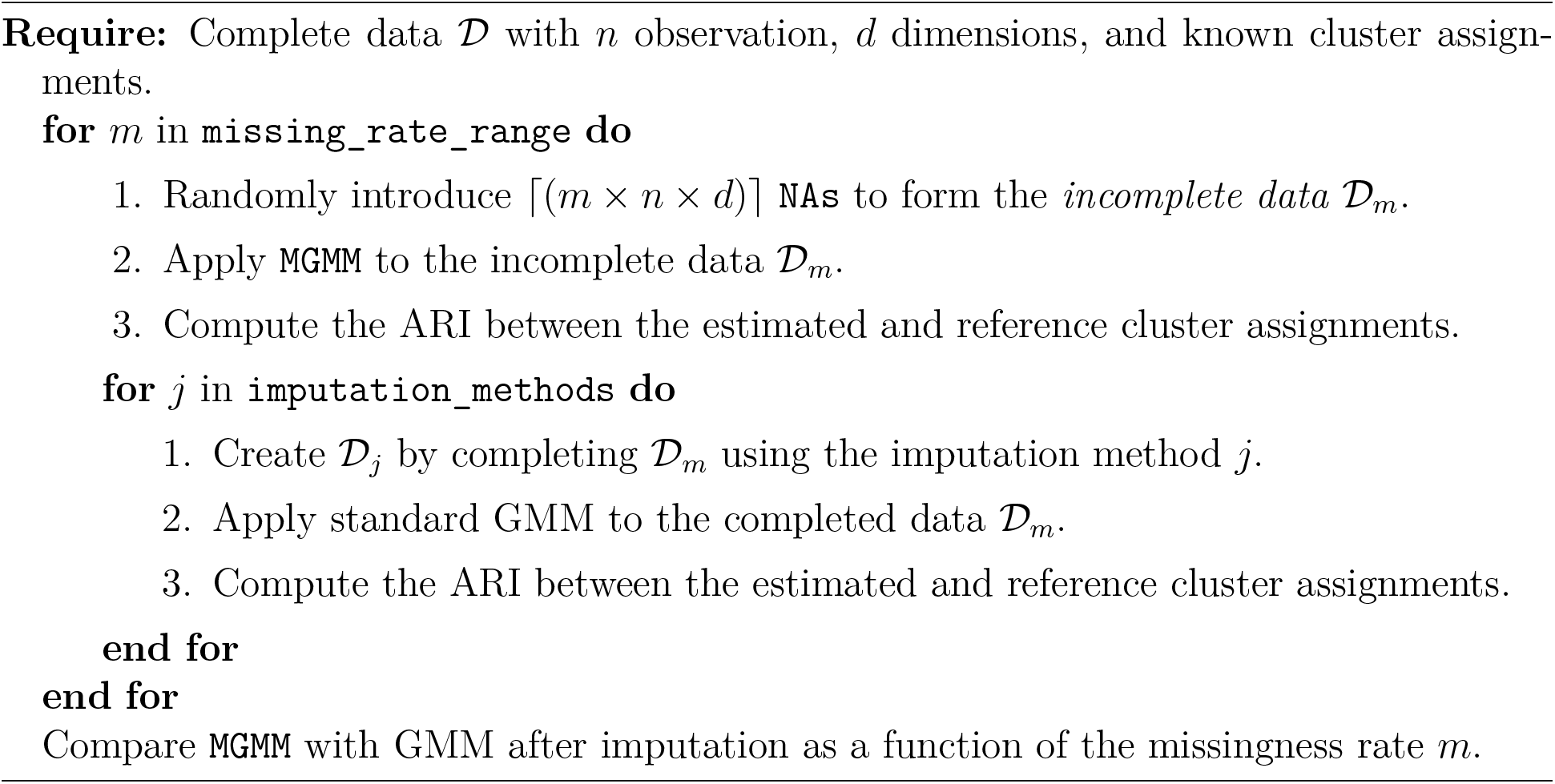
Benchmarking Procedure for `MGMM`

### 4.2 Benchmark Data Sets

#### 4.2.1 Simulated Gaussian Mixture

For the first clustering task, we consider data that were truly generated from a GMM, which is the setting in which `MGMM` should perform optimally. Data were simulated according to the hierarchical model in (2.1). The dimensionality *d* was set to 2 and the number of cluster components *k* to 4. The marginal density of the data generating process was:

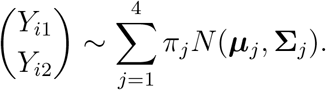

The means (***μ***_*j*_) were drawn from a uniform distribution on the square:

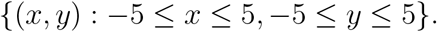

The component variances were set to 0.9:

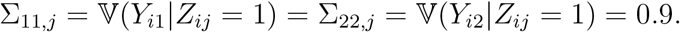

The covariance was uniformly sampled from the interval (−0.9, 0.9):

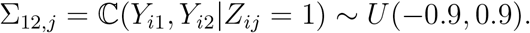

Marginal membership to each cluster was equally likely *π_j_* = 0.25 for *j* ∈ {1,⋯, 4}. A sample of size *n* = 2000 was generated using the `rGMM` function. The true (generative) component memberships (***z***_*i*_) were used as the reference when evaluating clustering performance on incomplete data.

#### 4.2.2 RNA Sequence Data from Cancer Patients

For the second clustering task, cancer gene expression data (Weinstein et al., 2013) were retrieved from the University of California Irvine machine learning repository. These data consist of expression values for 20,531 genes from *n* = 801 patients having 1 of *k* = 5 tumor types. Marginal analysis of variance was performed to identify the 20 most significantly differentially expressed genes (DEGs) across tumor types. The patient by DEG matrix was then decomposed via principal components analysis (PCA), and the final cluster task was performed on the *d* = 5 leading principal components. *Y_il_* represents the expression of patient *i* along the *l*th principal component. The patient’s observed tumor type was used as the reference when evaluating clustering performance on incomplete data. The tumor types are abbreviated as follows:

- BRCA: Breast carcinoma.
- COAD: Colon adenocarcinoma.
- KIRC: Kidney renal clear-cell carcinoma.
- PRAD: Prostate adenocarcinoma.
- LUAD: Lung adenocarcinoma.

#### 4.2.3 GWAS Summary Statistics

For the third clustering task, we consider summary statistics, both simulated and real, arising from GWAS for cardiovascular disease risk factors. In this setting, *i* indexes single nucleotide polymorphisms (SNPs) and *Y_il_* is the standardized score (i.e. Z-score) quantifying the magnitude of the observed association between SNP *i* and phenotype *l*. The SNPs may belong to one of *k* clusters, where the Z-scores of SNPs within a cluster may exhibit correlations due to the combination of environmentally-induced correlation of the traits and sample overlap between the GWAS in which the Z-scores were ascertained.

A simulated set of GWAS summary statistics was generated for *d* =3 traits and 900 SNPs arising from 1 of *k* = 3 clusters. The marginal density was:

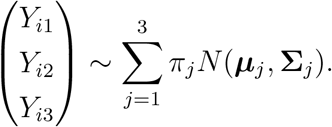

The mean vectors (***μ***_*j*_) were set to zero, and the cluster covariances (**Σ**_*j*_) were set to:

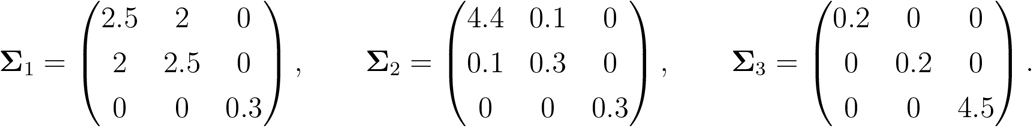

Marginal membership to each cluster was equally likely *π_j_* = 0.33 for *j* ∈ {1,⋯, 3}. A sample of size *n* = 900 was generated using the `rGMM` function from the `MGMM` package. To emulate the omission of non-significant results, which frequently occurs when reporting GWAS summary statistics, the SNPs were filtered to those having evidence of association with the traits at *p* ≤ 0.05 via the omnibus test (12) detailed in the appendix. After filtering, *n* = 183 SNPs remained, with marginal cluster frequencies: *π*_1_ = 0.25, *π*_2_ = 0.42, *π*_3_ = 0.33. The topology of the resulting data set is presented in Figure 2. The true (generative) component memberships (***z***_*i*_) were used as the reference when evaluating clustering performance on incomplete data.

**Figure 2:**
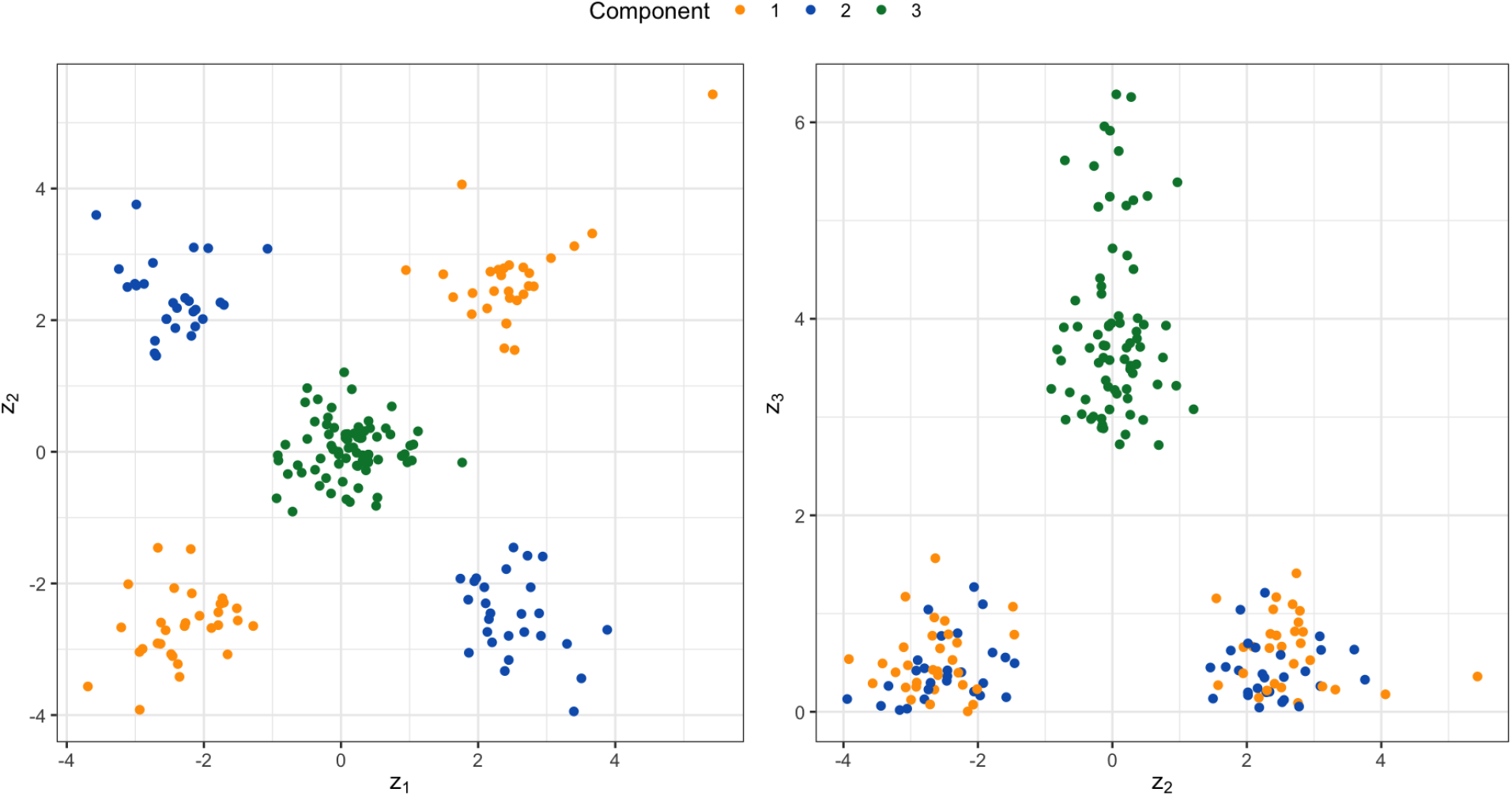
Scatter Plot of the Simulated GWAS-like Multivariate Z-scores. Observations are colored according to component membership. The left panel plots the second coordinate against the first, and the right panel plots the third coordinate against the second.

A set of real GWAS summary statistics for cardiovascular disease risk factors was prepared as described in Julienne et al. (2020b). These traits were: body mass index (BMI), coronary artery disease (CAD), low density lipoprotein (LDL), triglycerides (TG), waist to hip ratio (WHR), and any strokes (AS). From this collection of traits, we formed three example data sets. The first included {BMI, CAD, LDL} only; the second included {LDL, TG, BMI, AS, CAD}; the third {LDL, TG, BMI, AS, WHR}. We selected independent SNPs that were genome-wide significant (p-value ≤ 10^-8^) either marginally or via the omnibus test (12). These data sets contained 165, 166 and 179 SNPs respectively. For each example, a GMM with *k* = 3 components was fit to the complete data (using `fit.GMM` from the `MGMM` package), and the cluster assignments from this initial model were used as the reference when evaluating clustering performance on incomplete data. Because the reference clustering partition was directly derived from the data, the benchmark on these examples assess the robustness of the clustering rather than the ability to recover a true, underlying data class assignment.

### 4.3 Imputation Methods

In the absence of a missingness-aware method for fitting GMMs, the input data must either be reduced to complete cases, or completed via imputation. We consider the following imputation strategies: naive mean and median imputation, k-nearest neighbor (kNN) imputation Kowarik and Templ (2016), multiple imputation by chained equations (MICE) Buuren and Groothuis-Oudshoorn (2010), and random forest imputation Stekhoven and Bühlmann (2011). Naive mean or median imputation refers to simply setting a missing value to the mean or median of the observed values along that coordinate. For kNN, a missing value was imputed to the (Euclidean) distance-weighted average of the 5 nearest observations with observed data along that coordinate. For MICE, a missing value was imputed to its conditional expectation given the observed coordinates via the method of predictive mean matching; the number of imputations was 10, and the maximum number of Gibbs sampling iterations was 50. For random forest imputation, the number of trees per forest was 100, and the maximum number of refinement iterations was 10.

### 4.4 Benchmark Results

#### 4.4.1 Four Component Mixture of Bivariate Gaussians

When the underlying distribution was in fact a GMM, MGMM uniformly dominated imputation plus GMM (Figure 3) at recovering the true cluster assignments. Imputation by kNN and MICE performed similarly. Interestingly, although non-parametric, random forest imputation was not competitive when the true data generating process was a GMM. Naive mean and median imputation strongly under-performed, and at elevated missingness created singularities in the data set that prevented the GMM from converging.

**Figure 3:**
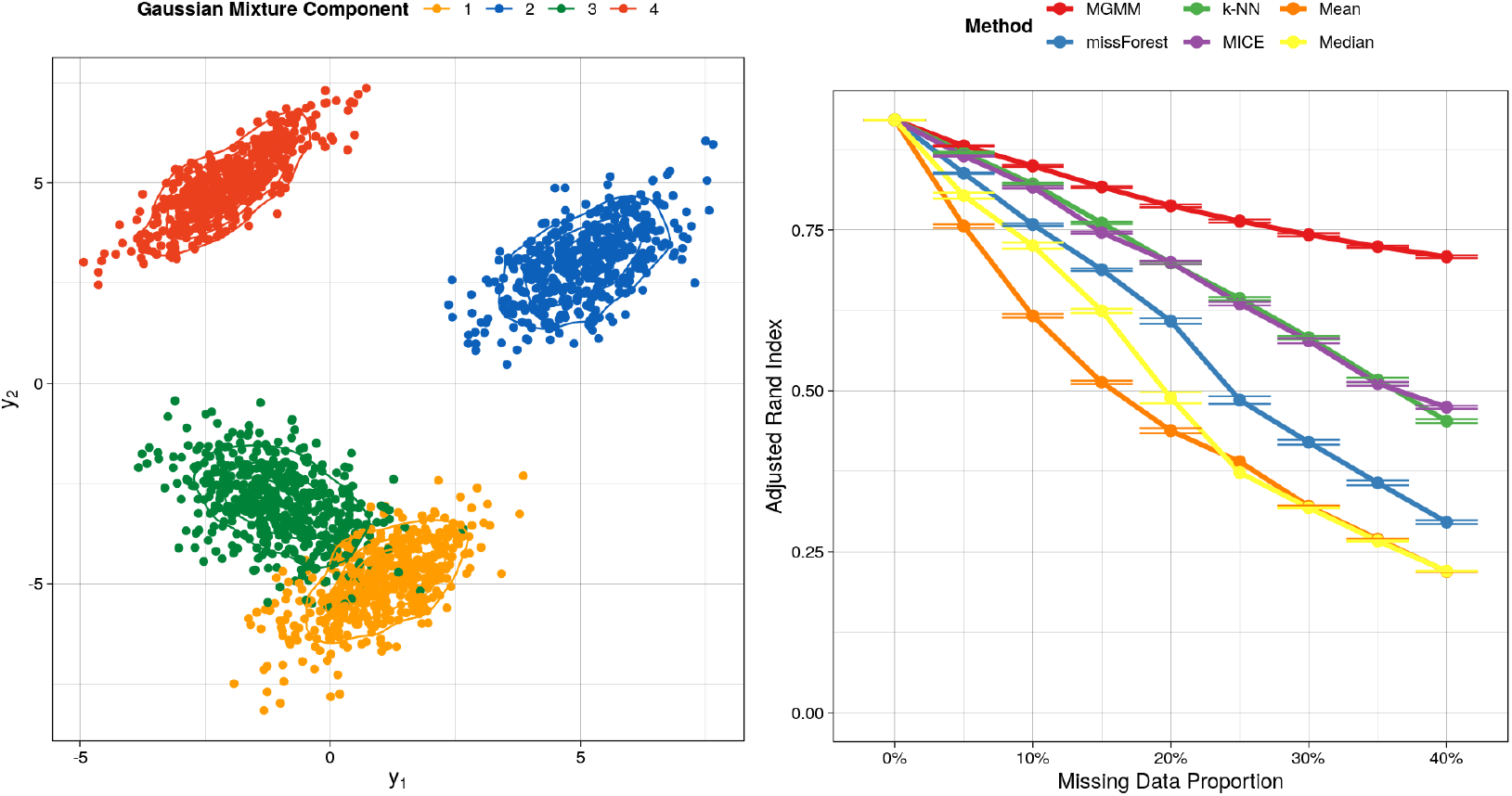
Benchmarking for the Mixture of Gaussians Data Set. The left panel includes the observations as simulated, colored according to the mixture component. The right panel presents the ajusted Rand index as a function of the missing data proportion for 8 different approaches to handling missing data; a higher value indicates better agreement between the predicted and true cluster assignments, adjusting for chance. Error bars represent the standard error of the mean across 20 simulation replicates.

#### 4.4.2 RNA Sequence Data from Cancer Patients

For the Cancer RNA-Seq data set, where the true generative model is unlikely to be a GMM, MGMM remained highly effective at recovering the true tumor type of the patient (see Figure 4). Random forests and kNN were competitive with MGMM, and outperformed when the proportion of missing data was ≥ 35%. Mean and median imputation were again not competitive, particularly when the proportion of missing data was ≥ 20%. MICE performed only slightly better than mean and median imputations. Linear imputation method may be ill-suited for separating the BRCA, LUDA, and PRAD tumor types.

**Figure 4:**
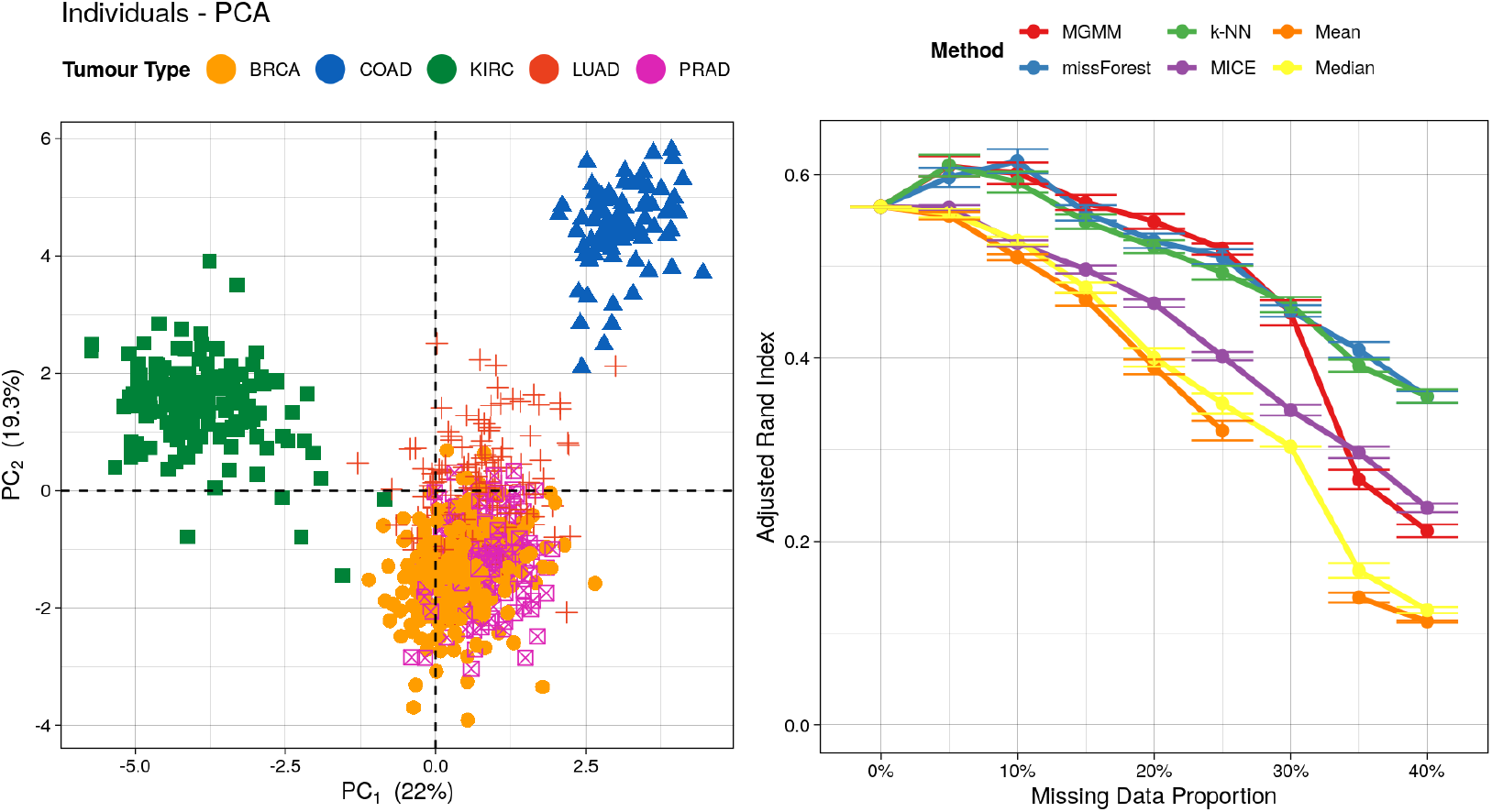
Benchmarking for the Cancer RNA-Seq Data Set. The left panel includes the projection of the expression data for *n* = 801 cancer patients onto the first two principal components. Observations are colored according to tumor type. The right panel presents the adjusted Rand index as a function of the missing data proportion; a higher value indicates better agreement between the predicted and true cluster assignments, adjusting for chance. Error bars represent the standard error of the mean across 20 simulation replicates.

#### 4.4.3 GWAS Summary Statistics

Finally, we considered clustering vectors of GWAS summary statistics arising when the same SNPs are tested for association with multiple traits. This analysis is of interest for identifying pleiotropy, individual SNPs that have effects on multiple traits, and polygenicity, collections of multiple SNPs that have effects on common traits. Such analyses are often performed by combining data from multiple independent studies, and missingness arises because not all SNPs or all traits were ascertained in all studies. Further, this analysis would generally only include SNPs that were significantly associated with at least one trait.

Here we discuss one simulated and one real data example; two additional real data examples are presented in the appendix. For the simulated summary statistics in Figure 5, the clustering task is same as the one presented on Figure 3. The three clusters are clearly separated. Yet, the task remains challenging due to the specific and unusual cluster topology arising from GWAS data. Since the underlying distribution was in fact a GMM, MGMM again performs very well, only falling off when the proportion of missing data reaches 40%. As for the RNA-Seq example (Figure 4), random forest and kNN were competitive with MGMM, outperforming at very high missingness. Surprisingly, MICE under-performed naive mean imputation, and was comparable to native median imputation.

**Figure 5:**
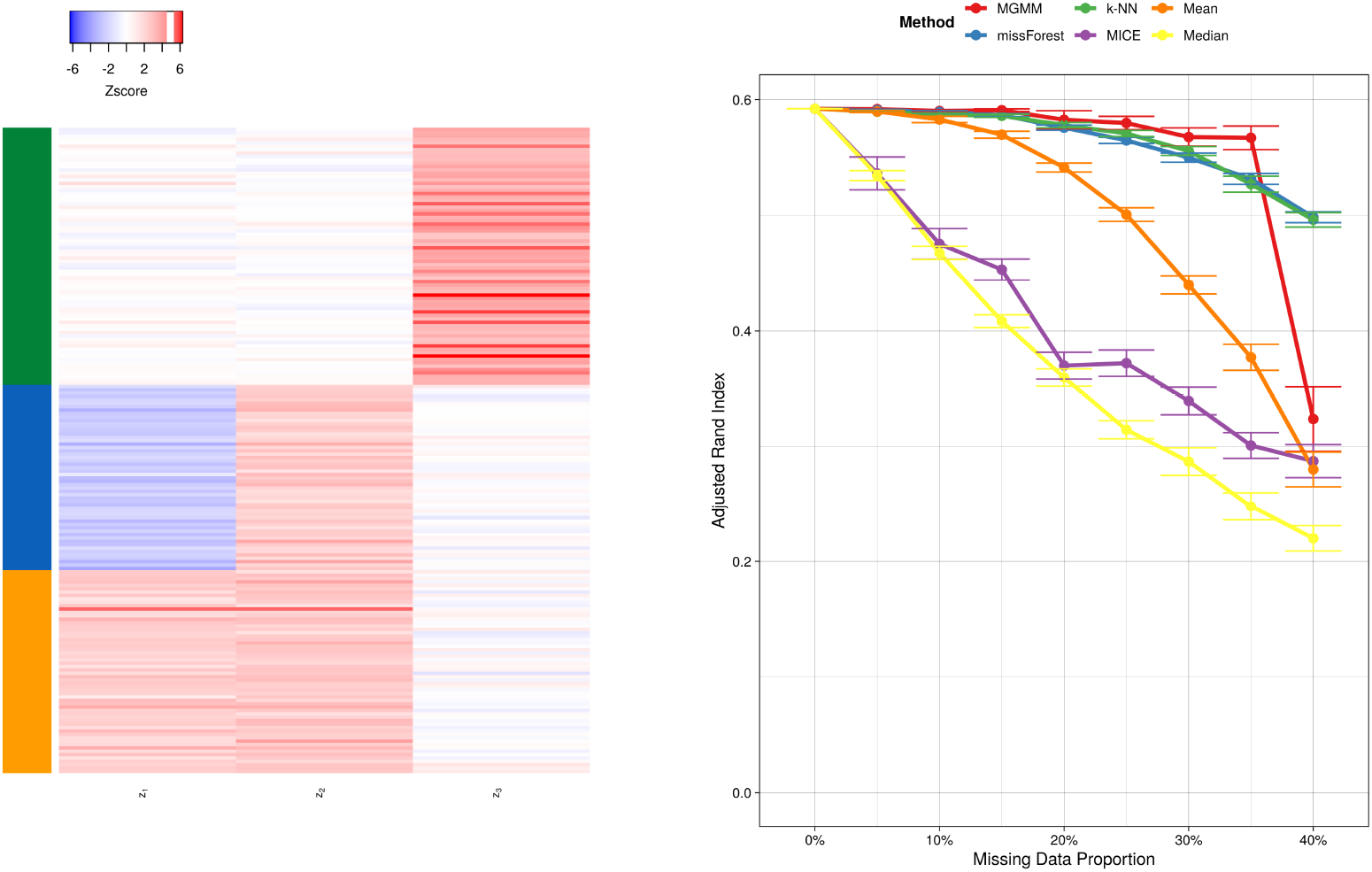
Benchmarking Simulated Multi-trait GWAS Summary Statistics. The left panel presents a heat map colored according to the normalized genetic effect, with SNPs as rows and traits as columns. The colorbar on the left represents the true cluster assignments. The right panel presents the adjusted Rand index as a function of the missing data proportion; a higher value indicates better agreement between the predicted and true cluster assignments, adjusting for chance. Error bars represent the standard error of the mean across 20 simulation replicates.

An analogous clustering task applied to summary statistics from real GWAS of BMI, CAD, and LDL is presented in Figure 6. The three clusters, identified by applying a 3-component GMM to the data before the introduction of missingness, appear well-differentiated on the heat map. kNN and random forests offered the best performance, followed by MGMM, whose performance deteriorated at missingness ≥ 35%. The deficit in performance of MGMM compared to kNN and random forests, even at low missingness, likely reflects a departure of the true data generating process from a GMM. As in the case of simulated GWAS summary statistics, MICE was not competitive, performing similarly to naive mean and median imputation.

**Figure 6:**
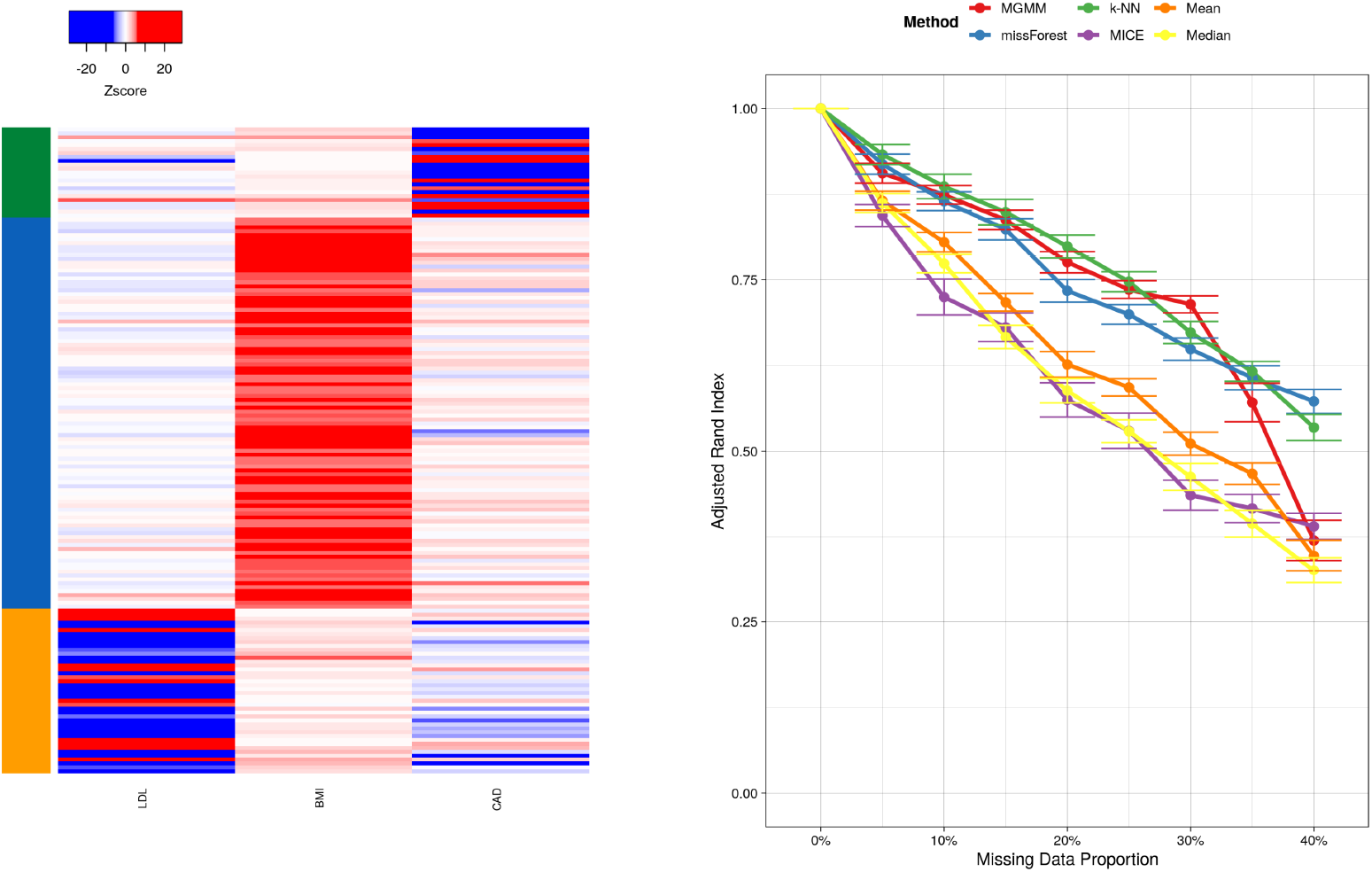
Benchmarking Real Multi-trait GWAS Summary Statistics for 3 Cardiovascular Risk Factors. These were body mass index (BMI), coronary artery disease (CAD), and low density lipoprotein (LDL). The left panel presents a heat map colored according to the standardized genetic effect, with SNPs as rows and traits as columns. The colorbar on the left represents the true cluster assignments. The right panel presents the adjusted Rand index as a function of the missing data proportion; a higher value indicates better agreement between the predicted and true cluster assignments, adjusting for chance. Error bars represent the standard error of the mean across 20 simulation replicates.

## 5 Filtering Unassignable Observations from MGMM and MICE

Since both MGMM and MICE provide an indication of the uncertainty in the cluster assignments, we created an additional clustering method in which observations with high assignment uncertainty were regarded as unassignable. This occurs when an observation could very plausibly has originated from more than one of the clusters, and may be exacerbated by excess missing data along a coordinate that helps to differentiate the clusters.

This uncertainty can be assessed via the entropy of the posterior membership probabilities:

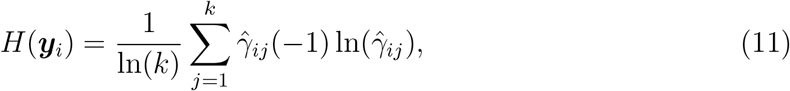

where 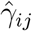 is the final responsibility of cluster *j* for observation *i*.

For MGMM, the entropy of the posterior cluster responsibilities is calculated by `fit.GMM` via (11). For MICE, each input data set is multiply imputed, and each of these imputed data set results in one *maximum a posteriori* cluster assignment. The posterior probability of membership to each cluster (i.e. the responsibilities, 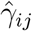) may be approximated by the proportion of imputations on which an observation was assigned to each cluster.

### 5.1 Filtering Procedure

In the filtered versions of MGMM and MICE, observations with high assignment uncertainty are identified via entropy and removed from consideration. For a given data set, such as the Cancer RNA-Seq data set, the distribution of entropy for MICE was typically right skewed (Figure 7-A). Consequently, for a fixed entropy threshold, the fraction of observations deemed unassignable is systematically higher for MICE than for MGMM (see Figure 7-A). To conduct a fair comparison of the two methods, we proceeded as follows:

1. For MICE, filter out observations with entropy exceeding 0.2 and assess performance on the remaining data.
2. Find the fraction of observations discarded by MICE *f*_disc_.
3. Determine the upper *f*_disc_th quantile *q*_MGMM_ of MGMM entropy. That is, find the threshold *q*_MGMM_ such that the proportion of observations with MGMM entropy exceeding *q*_MGMM_ is *f*_disc_.
4. For MGMM, filter out observations with entropy exceeding *q*_MGMM_ and assess performance on the remaining data.

**Figure 7:**
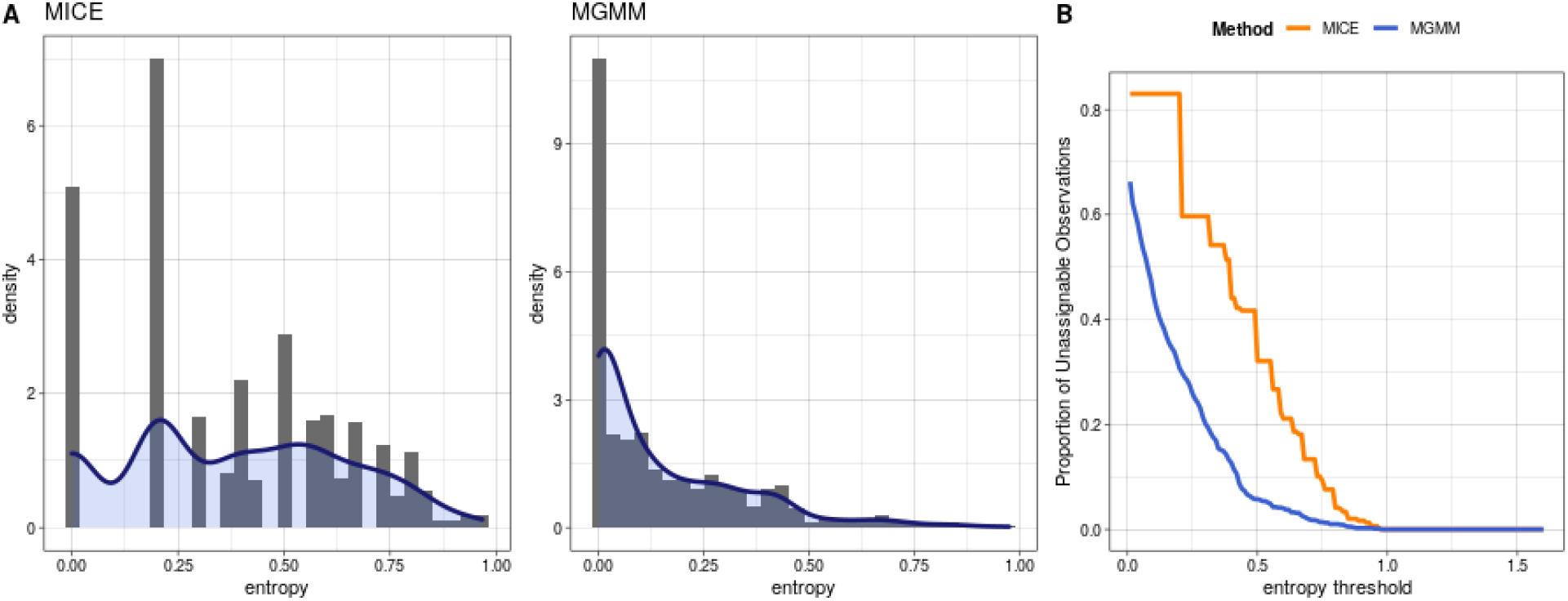
Comparison of Entropy Distributions of MICE and MGMM. A) Entropy distribution for each method on the Cancer RNA-Seq Data Set at a missing value ratio of 20%. B) Fraction of the data set deemed unassignable as a function of the entropy threshold.

This procedure provides a fair comparison of MGMM-filtered and MICE-filtered by adaptively selecting the entropy threshold for MGMM in such a way that both methods remove the same fraction of high uncertainty observations.

### 5.2 Comparison of MICE-filtered and MGMM-filtered

By effectively removing poorly classifiable observations from consideration, filtering is expected to improve the clustering quality, but only if those observations with high assignment uncertainty are correctly identified. Therefore, the comparative performance of MGMM-filtered and MICE-filtered provides an indication of how well each strategy was able to identify those observations with high cluster assignment uncertainty. We present the performances of the two methods on four data sets in Figure 8

**Figure 8:**
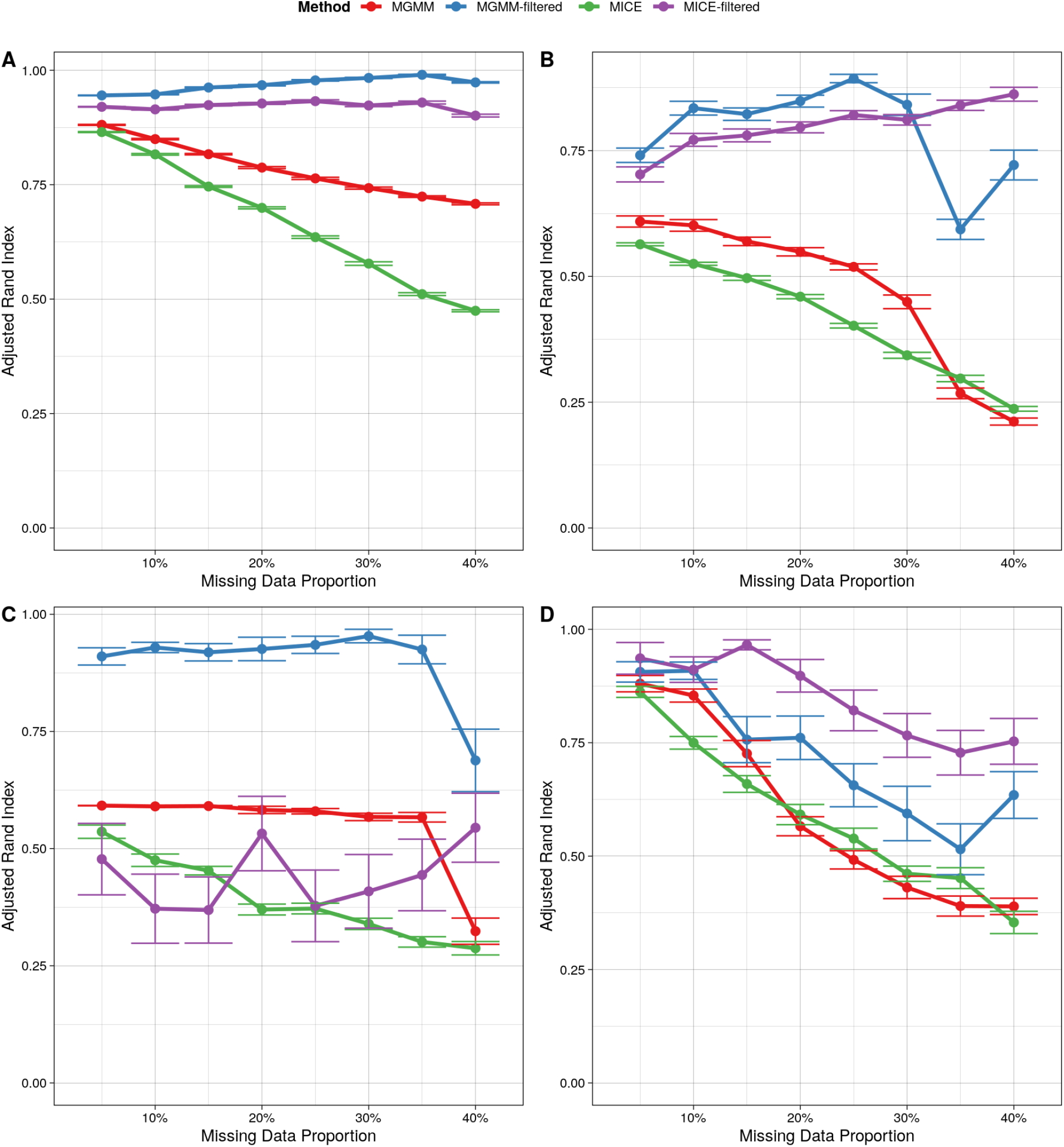
Performances of MICE-filtered and MGMM-filtered on four Benchmark Data Sets. The adjusted Rand index as a function of the missing data proportion for A) the Four Component Mixture of Bivariate Gaussians simulation, B) Cancer RNA-Seq Data Set, C) Simulated Multi-trait GWAS Summary Statistics, D) 2nd Example of Real Multitrait GWAS Summary Statistics for Cardiovascular Risk Factors. Error bars represent the standard error of the mean across 40 simulation replicates.

#### 5.2.1 Four Component Mixture of Bivariate Gaussians

For the Gaussian mixture simulation data set (Figure 8A), filtering out unassignable observations strikingly improved the classification accuracy of both MICE and MGMM. However, MGMM-filtered performed better for all missing data ratios. Thus, when the data are in fact generated by a GMM, MGMM correctly assesses cluster assignment uncertainty, providing users with a mechanism for identifying observations with low-confidence cluster assignments.

#### 5.2.2 RNA Sequence Data from Cancer Patients

For the cancer RNA-Seq data set (Figure 8B), entropy-based filtering again significantly improved the performance of both methods, suggesting that assignment entropy provides a reliable method for identifying unassignable observations. Note that the filtered data set contained sufficiently many observations to correctly evaluate performance (see Appendix 9). MGMM-filtered outperformed MICE-filtered at lower missingness, while MICE-filtered performed better at a missing data ratio exceeding 30%. The same trend was observed for the unfiltered versions of MGMM and MICE. This example demonstrates that even when the underlying distribution is not a GMM, MGMM is able to accurately assess cluster assignment uncertainty at practical missing ratios.

#### 5.2.3 GWAS Summary Statistics

For the GWAS summary statistic data sets, the comparative performances of the two methods depend on the structure of the data. On the simulated multi-trait GWAS summary statistics (Figure 8C), filtering drastically improved the performance of MGMM, whereas filtering did little, if anything, to improve the performance of MICE. This suggest that MICE-based imputation entropy was not an effective gauge of assignment uncertainty for these data. The non-linearity and absence of correlation among the variables probably explains the poor performance of MICE.

On the 2nd example of real GWAS summary statistics for cardiovascular risk factors, MICE-filtered performed best overall, and entropy-based filtering improved the performance of MICE more so than the performance of MGMM. The unfiltered versions of MICE and MGMM performed comparably. The strong correlations among the traits studied likely explains the good performance of MICE for these data. It is also important to note that, for the GWAS data sets, the reference labels used to compute the adjusted rand index are not the true classes *per se*, but rather the clustering obtained on complete data (see 4.2.3). Therefore, the performance assessment in this example is more a measure of the robustness of the clustering procedure to the missing data ratio than a measure of the capacity to identify true underlying classes.

#### 5.2.4 Filtered Observations

Importantly, filtering out unassignable observations based on entropy did not strongly enrich the remaining data for complete cases (see Figure 12). Therefore, the general improvements in performance observed with filtering cannot be trivially explained by the selective removal of incomplete observations, and point instead to the accurate identification of observations that could plausibly have arisen from more than 1 cluster.

## 6 Conclusion

We have presented `MGMM`, a powerful, general purpose `R` package for maximum likelihood based estimation of GMMs in the presence of missing data, and demonstrated that MGMM often outperforms imputation followed by standard GMM on various real and simulated data sets. In contrast to imputation, `MGMM` uses the ECM algorithm to efficiently obtain unbiased estimates of model parameters by maximizing a lower bound on the observed data log likelihood. The functionalities of the `MGMM` package are carefully documented and comprise: the generation of random data under a specified GMM, the fitting of GMM parameters to data sets containing missing values, and the computation of a panel of clustering criteria to identify the optimal number of clusters. We conducted a comparative benchmark to assess the capacity of MGMM to correctly identify true cluster assignments in data containing missing values as compared with imputation followed by a classical GMM. We established that for data sets following a distribution close to a GMM, MGMM is able to recover the true class assignment more accurately than imputation followed by a classical GMM. When the underlying data generating process is in fact a GMM, then as a correctly specified maximum likelihood procedure, MGMM is optimal. Moreover, MGMM correctly assess its level of uncertainty in clustering assignments, providing a mechanism for identifying and separating out unassignable observations.

GMMs are not well-suited to all clustering tasks. Direct application of MGMM was less effective than non-linear imputation, via kNN or random forests, followed by standard GMM in cases where the clusters present in the observed data were poorly differentiated, or the missingness was high (e.g. 40%). This observation emphasizes the need to assess the appropriateness of a GMM before applying MGMM to a clustering problem. Since kNN and random forest imputation, followed by standard GMM, were typically competitive with MGMM in the real data examples, these methods may be used to perform sensitivity analysis on the final cluster assignments. On the other hand, standard GMM following kNN or random forest imputation will not appropriately propagate uncertainty due to missing data. This can lead to inaccurate estimates of the posterior membership probabilities, particularly for observations with multiple missing elements, and failure to identify observations whose cluster assignments are unreliable. Thus, an approach such as MGMM-filtered, which accurately assesses assignment uncertainty and removes unclassifiable observations from consideration, may be more reliable. The framework proposed by Ghahramani and Jordan (1994) and elaborated upon here, of using an EM-type algorithm to fit mixture models in the presence of both missing data and unknown class assignments, may be extended to estimates mixtures of non-Gaussian distributions. Extending MGMM to estimate such mixtures in the presence of missing data is among our future directions.

## Appendix

### 7 Omnibus Association Test

The *omnibus test* is a classic, multi-trait test for assessing the association between a given SNP and any of a set of traits. Let **z** denote a *d* dimensional vector of Z-scores quantifying the association between the SNP and the d traits. These are (asymptotically) normally distributed, with mean ***μ*** and covariance **Σ**:

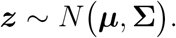

Note that covariance arises among the Z scores when the traits are correlated. The null hypothesis of the omnibus test is that the means of all summary statistics are zero:

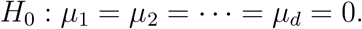

The alternative is that at least one mean is non-zero. The omnibus test statistic is the quadratic form:

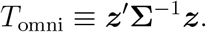

Under the null, *T*_omni_ follows a central 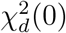 distribution with *d* degrees of freedom:

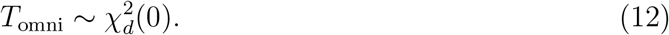

### 8 Additional Analyses of GWAS Summary Statistics

Here we present two additional examples of clustering summary statistics from real GWAS of cardiovascular disease risk factors. In both cases, the underlying data generating mechanism is unknown, but is unlikely to be a GMM. One cluster (green), corresponding to SNPs associated with BMI, is well differentiated, while the remaining two clusters (blue and gold) are poorly resolved, and it is not obvious in either case that there truly are thee clusters present in the observed data. In this context, non-linear imputation, via kNN or random forests, followed by standard GMM outperformed MGMM. This is not surprising: when the observed data arise from a distribution that is far from a GMM, using MGMM to circumvent imputation is unlikely to succeed. These examples underscore the point made in the conclusions that GMMs, and MGMM in particular, are not suited to all clustering tasks.

**Figure 9:**
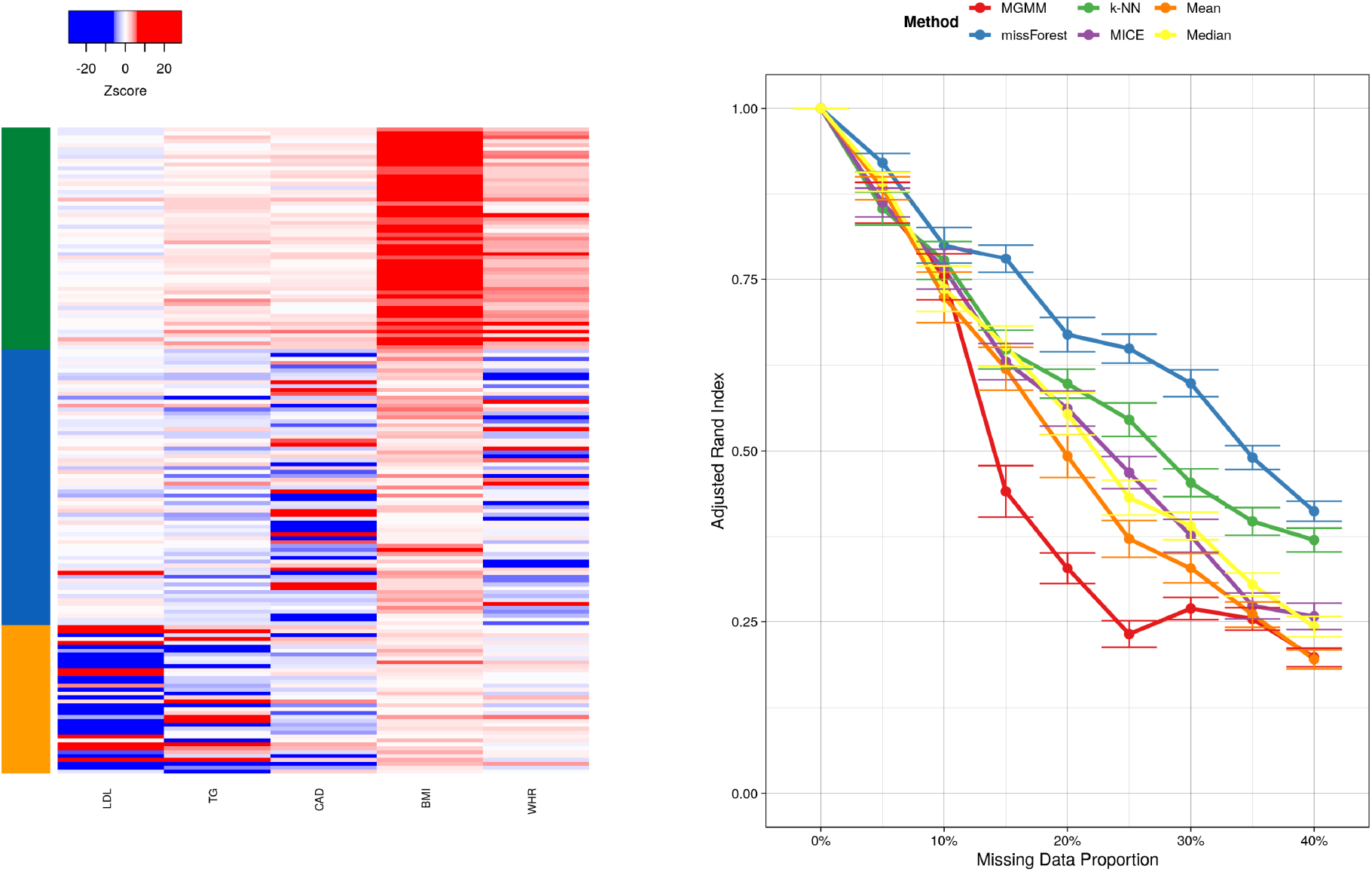
Benchmarking Real Multi-trait GWAS Summary Statistics for 5 Cardiovascular Risk Factors. These were: body mass index (BMI), coronary artery disease (CAD), low density lipoprotein (LDL), triglycerides (TG), and waist-to-hip ratio (WHR). The left panel presents a heat map colored according to the standardized genetic effect, with SNPs as rows and traits as columns. The colorbar on the left represents the true cluster assignments. The right panel presents the adjusted Rand index as a function of the missing data proportion; a higher value indicates better agreement between the predicted and true cluster assignments, adjusting for chance. Error bars represent the standard error of the mean across 20 simulation replicates.

**Figure 10:**
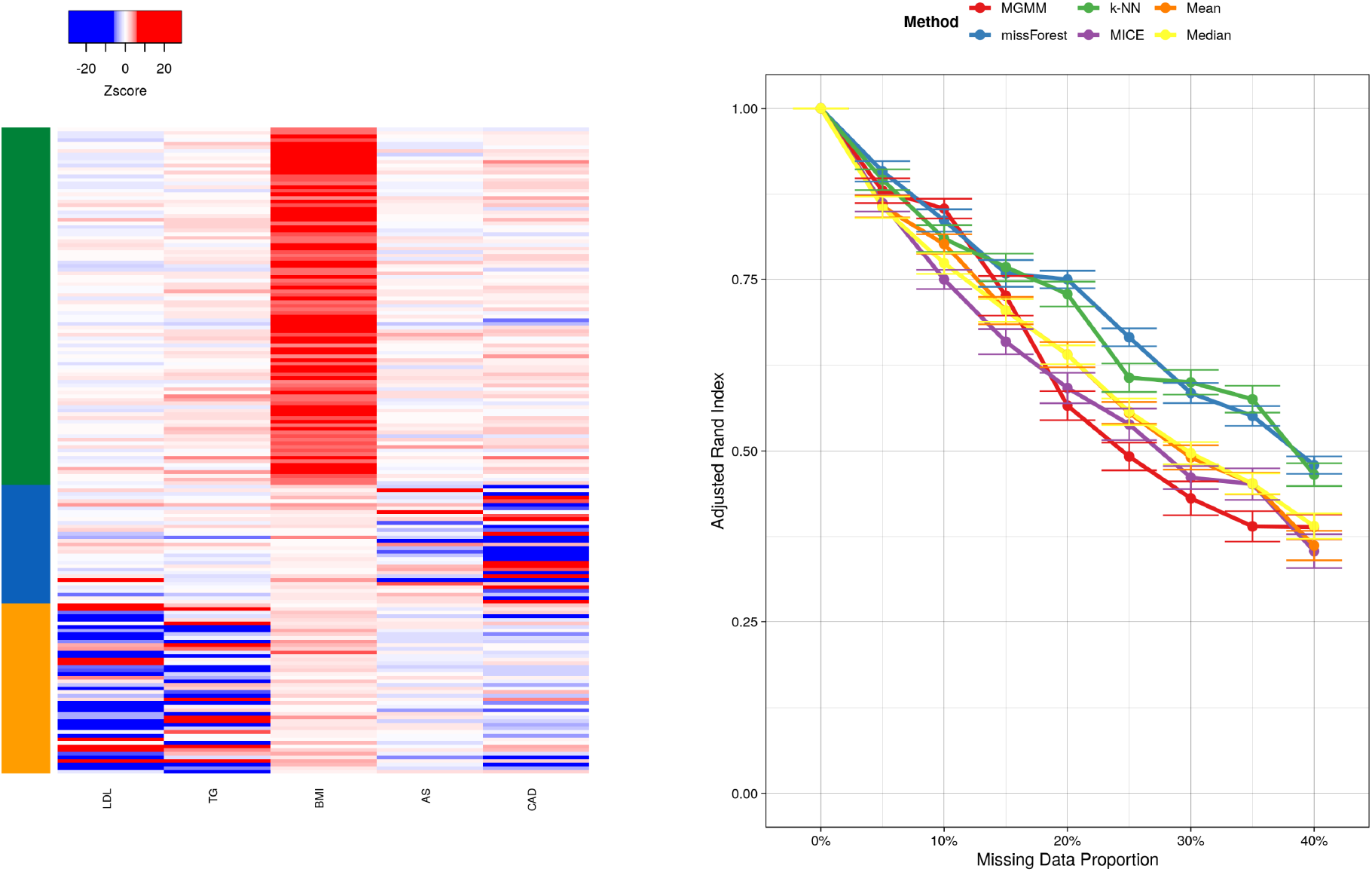
2nd Example of Benchmarking Real Multi-trait GWAS Summary Statistics for 5 Cardiovascular Risk Factors. These were: any strokes (AS), body mass index (BMI), coronary artery disease (CAD), low density lipoprotein (LDL), triglycerides (TG). The left panel presents a heat map colored according to the normalized genetic effect, with SNPs as rows and traits as columns. The colorbar on the left represents the true cluster assignments. The right panel presents the adjusted Rand index as a function of the missing data proportion; a higher value indicates better agreement between the predicted and true cluster assignments, adjusting for chance. Error bars represent the standard error of the mean across 20 simulation replicates.

### 9 Comparison of the MICE-filtered and MGMM-filtered Procedures

We compared the fraction of observations deemed unassignable by the MICE-filtered and MGMM-filtered to ensure that filtering did not lead to an over-enrichment of complete observations, as this might have artificially increased the apparent performance. Figure 11 presents the fraction of observations deemed unassignable from the cancer RNA-Seq data set, and the second 5-trait real GWAS benchmark. Note that, by design of the filtering procedure, both methods remove the same proportion of observations, with minor variations due to the imprecision of the empirical distribution function for assignment entropy. As the missing data ratio increased, so too did the proportion of unassignable observations, likely due to a loss of information along coordinates important for differentiating the clusters.

**Figure 11:**
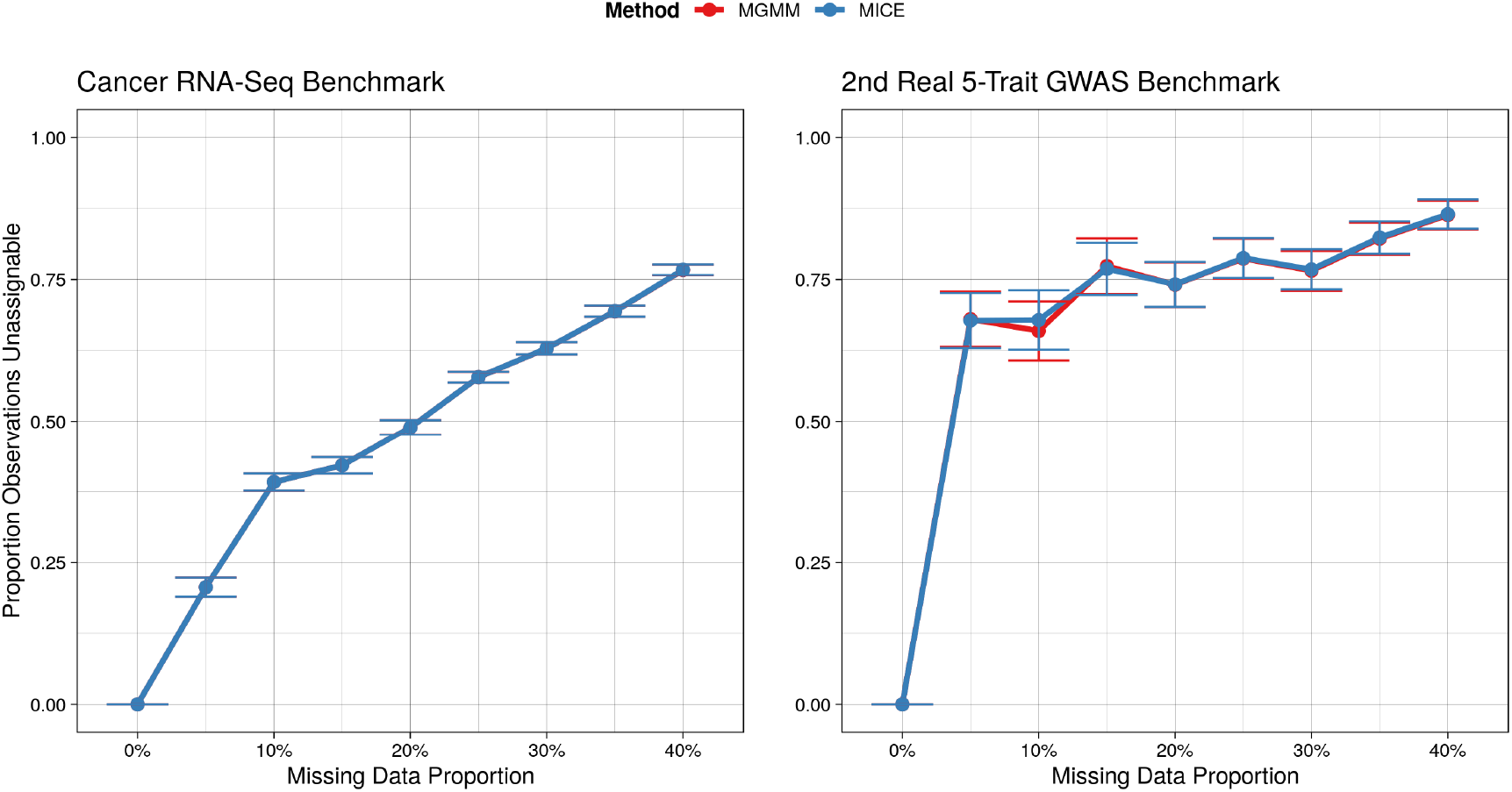
Proportion of Observations Deemed Unassignable as a Function of the Proportion of Missing Data. By design of the filtering procedure, both MGMM-filtered and MICE-filtered should remove nearly the same proportion of observations.

Figure 12 verifies that the proportion of complete observations remaining after filtering was not dramatically increased by either MGMM or MICE. The black dashed line on each panel represents the expected fraction of complete observations for a given missingness, assuming the missingness occurs completely at random. For missingness *m* and data of dimension *d*, this fraction is *f*_complete_ = (1 − *m*)^*d*^.

**Figure 12:**
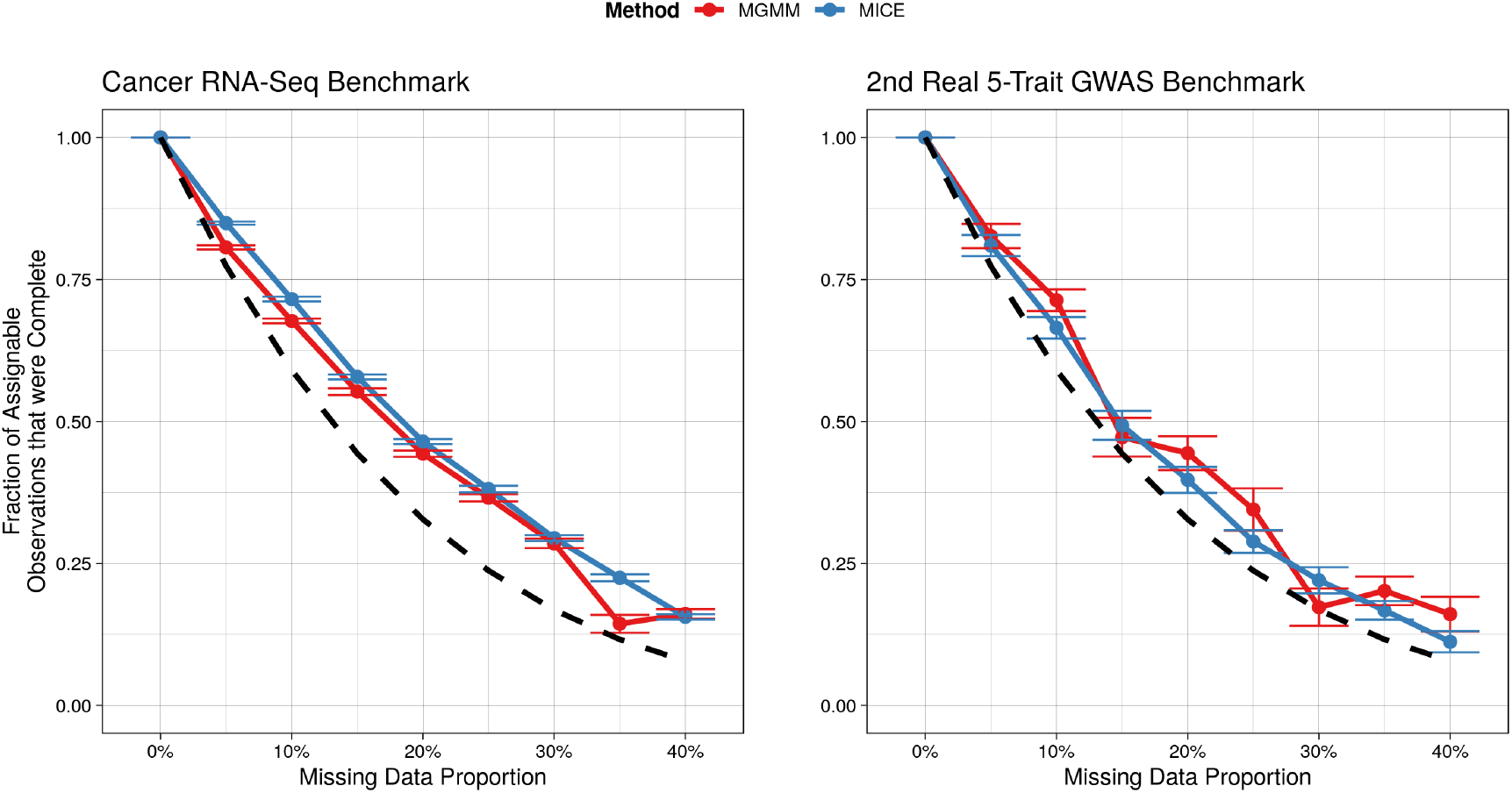
Proportion of Complete Observations Remaining among the Assignable Observations. The expected fraction of complete observation is represented by the dashed black curve.

